# Omicron Spike confers enhanced infectivity and interferon resistance to SARS-CoV-2 in human nasal tissue

**DOI:** 10.1101/2023.05.06.539698

**Authors:** Guoli Shi, Tiansheng Li, Kin Kui Lai, Reed F. Johnson, Jonathan W Yewdell, Alex A Compton

**Author notes:** These authors contributed equally (Guoli Shi, Tiansheng Li).

## Abstract

Omicron emerged following COVID-19 vaccination campaigns, displaced previous SARS-CoV-2 variants of concern worldwide, and gave rise to lineages that continue to spread. Here, we show that Omicron exhibits increased infectivity in primary adult upper airway tissue relative to Delta. Using recombinant forms of SARS-CoV-2 and nasal epithelial cells cultured at the liquid-air interface, enhanced infectivity maps to the step of cellular entry and evolved recently through mutations unique to Omicron Spike. Unlike earlier variants of SARS-CoV-2, Omicron enters nasal cells independently of serine transmembrane proteases and instead relies upon metalloproteinases to catalyze membrane fusion. This entry pathway unlocked by Omicron Spike enables evasion of constitutive and interferon-induced antiviral factors that restrict SARS-CoV-2 entry following attachment. Therefore, the increased transmissibility exhibited by Omicron in humans may be attributed not only to its evasion of vaccine-elicited adaptive immunity, but also to its superior invasion of nasal epithelia and resistance to the cell-intrinsic barriers present therein.

## Introduction

Efficient infection of human cells by severe acute respiratory syndrome coronavirus 2 (SARS-CoV-2) requires the interaction of SARS-CoV-2 Spike with its receptor at the cell surface, angiotensin-converting enzyme 2 (ACE2)^1,2^. In addition, Spike has been described to bind to various cellular factors to promote coronavirus attachment to the cell surface. 78-kilodalton glucose-regulated protein, neuropilin-1, high-density lipoprotein scavenger receptor B type 1, CD209/CD299, Axl, sialic acid, and heparan sulfate have all been reported to interact with Spike and promote the virus entry process^3-9^. In general, it is thought that factors aiding virus attachment enable subsequent ACE2 binding. Engagement of ACE2 alters Spike conformation and facilitates its processing by cellular proteases, such as serine transmembrane proteases like TMPRSS2, matrix metalloproteinases (MMP), or a disintegrin and metalloproteinases (ADAM)^2,10-14^. Protease cleavage enables Spike to trigger fusion between viral and cellular membranes to complete cellular entry^15^.

The Omicron B.1.1.529.1 (BA.1) variant emerged in late 2021 and, as a result of increased transmissibility between humans^16,17^, quickly replaced the Delta variant of concern as the dominant form of SARS-CoV-2 circulating worldwide (https://www.cdc.gov/coronavirus/2019-ncov/variants/variant-classifications.html). Compared to previous SARS-CoV-2 variants of concern, Spike from BA.1 contained a plethora of unique mutations, and its descendants (including the BA.2, BA.4, BA.5, BQ.1, and XBB subvariants) contained additional mutations in Spike (https://covid.cdc.gov/covid-data-tracker/#datatracker-home). The functional characterization of Omicron has revealed that said Spike mutations, including those found in the receptor binding domain, allow for evasion of neutralizing antibodies induced by COVID-19 vaccination^18-26^. Furthermore, some studies have shown that Omicron Spike displays increased affinity for human ACE2 compared to the early D614G variant^23,27,28^, but this was not supported by others^29^. Therefore, it remains unclear whether antibody evasion and ACE2 affinity are the only factors explaining the improved transmissibility of Omicron subvariants.

It has also been reported that Omicron utilizes a distinct cellular entry pathway compared to ancestral or early forms of SARS-CoV-2^30-34^. Specifically, the proteolytic activation of Omicron Spike is less dependent on TMPRSS2 but may be more dependent on cathepsin activity in endolysosomes or metalloproteinase activity at the plasma membrane^11^. However, the precise characterization of the entry pathways used by Omicron in physiologically relevant primary cells is lacking, and whether they are sensitive to cell-intrinsic antiviral barriers has not been determined. The apparent loss of TMPRSS2 dependence may have consequences for the cellular tropism of Omicron in vivo. Some reports suggest that Omicron BA.1 exhibits a decreased propensity for lower respiratory tract (lung) infection, which may explain the decreased pathogenicity associated with Omicron infections^35-40^. Since the upper respiratory tract, and specifically the nasal epithelium, represents the initial site of SARS-CoV-2 infection and the initial protective innate immune response^41,42^, we compared the ability of Omicron to invade human nasal epithelia relative to ancestral SARS-CoV-2 and the preceding variant of concern, Delta, using a primary human nasal tissue culture model. We found that Omicron BA.1 and BA.2 exhibited a markedly enhanced infectivity in primary nasal epithelia compared to early SARS-CoV-2, and using recombinant virus, we show that Spike from Omicron, but not Delta, controlled this phenotype. Furthermore, despite Omicron eliciting a type-I interferon response in nasal tissue, Omicron Spike used a metalloproteinase-mediated entry pathway that was associated with evasion of interferon-inducible factors targeting virus entry. Collectively, these findings may explain the increased transmissibility of Omicron as well as its ability to displace more interferon-sensitive variants of SARS-CoV-2.

Overall, our findings suggest that the global spread of Omicron was fueled not only by increased ACE2 affinity and resistance to vaccine-induced adaptive immunity, but also by resistance to nasal cell-intrinsic entry inhibitors.

## Results

### Omicron BA.1 exhibits superior replicative fitness relative to early SARS-CoV-2 and Delta in primary human nasal epithelial cells

Primary human nasal epithelial cells isolated from three donors were pooled and cultured as submerged monolayers, which allows for the propagation of cells in the basal (undifferentiated) state. Immunofluorescence staining of acetylated tubulin indicated that mature cilia are lacking in cells cultured in this manner, although they express detectable levels of ACE2 **(Figure 1A)**. Nasal monolayers were inoculated with early/ancestral SARS-CoV-2 (USA-WA1/2020 strain, herein referred to as WA1) at a multiplicity of infection (MOI) of 0.05, and RT-qPCR of viral ORF1a RNA at multiple time points were suggestive of virus replication over time (**Figure 1B**). Productive infection was confirmed by demonstrating that cell culture supernatants contained infectious virus that could be titered on Vero E6-ACE2-TMPRSS2 cells by ORF1a qPCR and anti-nucleocapsid immunostaining (**Supplemental Figure 1A**). By comparison, inoculation of nasal monolayers with the same MOI of BA.1 resulted in far greater virus replication at all time points **(Figure 1B)**. In contrast, BA.1 did not replicate in submerged monolayers of human small airway (lung) epithelial cells (three pooled donors), while WA1 replicated with similar kinetics in nasal and lung cells **(Figure 1C)**. These data suggest that BA.1 displays a growth advantage specific to the upper airway of the human respiratory tract. BA.1 exhibited an approximately 100-fold greater replication capacity in nasal monolayers compared to WA1, either when measuring absolute **(Supplemental Figure 1B)** or relative **(Supplemental Figure 1C)** copy numbers of viral ORF1a. We found that this replication advantage by BA.1 was accompanied by an approximately 30-fold superior capacity to adhere to the nasal cell surface compared to WA1 **(Supplemental Figure 1D)**. We excluded that WA1 and BA.1 inoculants (initially titered on Vero E6-TMPRSS2 cells) contained different quantities of virus particles by inoculating HEK293T-ACE2 and Vero E6-TMPRSS2 cells in parallel. In contrast to our observations in nasal monolayers, WA1 and BA.1 infected these permissive cell lines to equal extents (**Supplemental Figure 1E and 1F**). Even when virus inoculants were calculated using an alternative virus titering strategy (based on viral ORF1a RT-qPCR instead of Vero E6-TMPRSS2 infection), BA.1 maintained a significant replicative advantage in nasal monolayers relative to WA1 (**Supplemental Figure 1G**). Therefore, BA.1 exhibits enhanced replicative potential in primary human nasal cells relative to WA1 and this is associated with improved cell surface adhesion by BA.1 Spike. Since BA.1 emerged in humans while the preceding variant of concern, Delta, was widespread, we compared the capacity for BA.1 and Delta to infect human nasal monolayers. In contrast to the orders of magnitude improvement exhibited by BA.1, Delta infected nasal cells only modestly better than WA1, and the difference was not statistically significant (**Figure 1D**). Therefore, the enhanced ability for BA.1 to infect primary human nasal epithelia emerged in the Omicron ancestor, not in Delta. Importantly, remdesivir, an inhibitor of the viral RNA polymerase, significantly reduced ORF1a levels in nasal cells inoculated with WA1, Delta, and BA.1, demonstrating that measurement of ORF1a by RT-qPCR is reflective of virus replication (**Figure 1E**).

**Figure 1.**
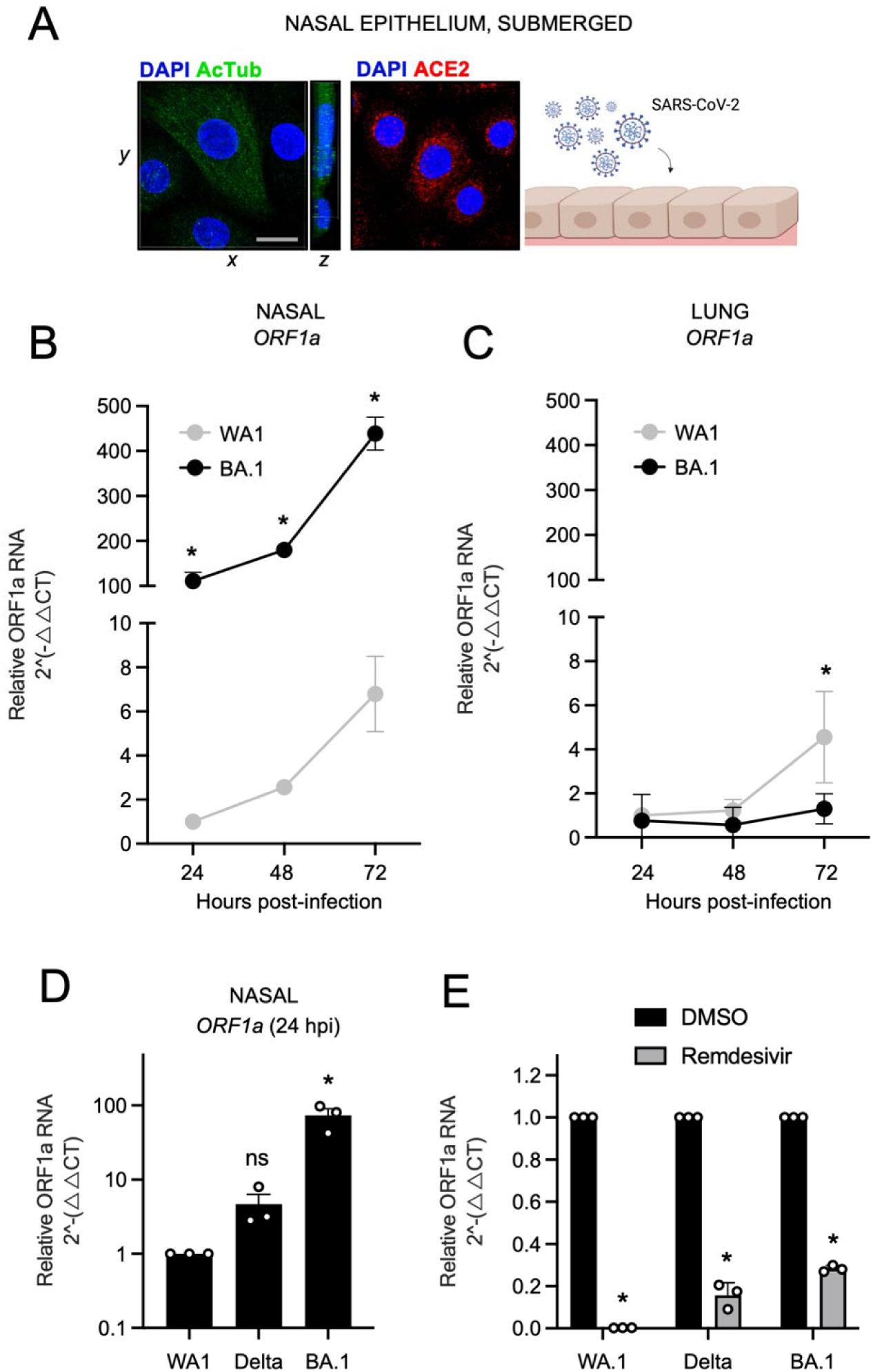
Omicron BA.1 exhibits superior replicative fitness relative to early SARS-CoV-2 and Delta in primary human nasal epithelial cells. (A) Primary human nasal epithelial cells (pooled from 3 human donors) were cultured as undifferentiated, submerged monolayers and challenged with SARS-CoV-2. Acetylated tubulin levels were determined by anti-AcTub immunofluorescence, ACE2 levels were determined by anti-ACE2 immunofluorescence, and DAPI was used to stain nuclei. 3D reconstructions of *xy* and *yz* fields are shown. Cartoon made with Biorender.com. (B) Primary human nasal epithelial cells (cells from three human donors, pooled) or (C) human small air way (lung) epithelial cells (cells from three human donors, pooled) were challenged with replication competent SARS-CoV-2 WA1 or Omicron BA.1 at an MOI of 0.05. Total cellular RNA was extracted and viral ORF1a was quantified by RT-qPCR at the indicated time points. Relative viral RNA abundance compared to actin was determined by the 2^(-ΔΔCT) method. ORF1a abundance of WA1 at 24 hours post inoculation was set to 1. (D) Virus replication in primary human nasal epithelial cells was measured by detecting ORF1a levels with RT-qPCR at 24 hours post inoculation with WA1, Delta, or BA.1 at an MOI of 0.05. (E) As in (D), except that primary human nasal epithelial cells were treated with remdesivir (10 µM) prior to and during inoculation with WA1, Delta, or BA.1. ORF1 abundance of each virus treated with DMSO was set to 1. All results are represented as means plus standard error from three independent infections (symbols represent biological replicates). Statistically significant differences (* *P* < 0.05) between the indicated condition and the corresponding data point of WA.1 were determined by one-way ANOVA.

During the preparation of this manuscript, it was reported that Omicron displays greater adhesion to human nasal cells, and this was attributed to enhanced binding to cilia in differentiated nasal epithelia^43^. Our results here suggest that, besides cilia, other factors intrinsic to nasal cells must govern the improved replicative fitness of BA.1 compared to early SARS-CoV-2.

### Omicron Spike enables enhanced SARS-CoV-2 infectivity in primary nasal epithelia cultured at the air-liquid interface

To recreate the three-dimensional, pseudostratified architecture of nasal epithelia in vivo, primary human nasal epithelial cells were pooled from 14 donors and differentiated at the air-liquid interface (ALI). In addition to ample ACE2 expression, differentiation status was confirmed by stratification of nuclei and the presence of mature cilia at the tissue surface **(Figure 2A)**. Relative to nasal monolayers (**Supplemental Figure 1B**), nasal ALI were more permissive to SARS-CoV-2 infection, as assessed by absolute ORF1a copy number (**Supplemental Figure 2A**). In this tissue, replication of Omicron BA.1 and BA.2 exceeded that of WA1 by multiple orders of magnitude at 48 hours post-inoculation, with BA.2 exhibiting the greatest replicative potential **(Figure 2B)**. To confirm that nasal ALI support repeated rounds of productive infection, we collected culture medium supernatants of nasal ALI inoculated with WA1, BA.1, or BA.2 and quantified infectious virus levels by titrating on Vero E6-TMPRSS2 cells. In agreement with RT-qPCR results, BA.1 and BA.2 achieved higher infectious titers in nasal ALI relative to WA1 **(Figure 2C)**. In accordance with enhanced virus replication of BA.1 and BA.2, we detected elevated type-I interferon induction (*IFNB*) at 48 hours post-inoculation **(Figure 2D)**.

**Figure 2.**
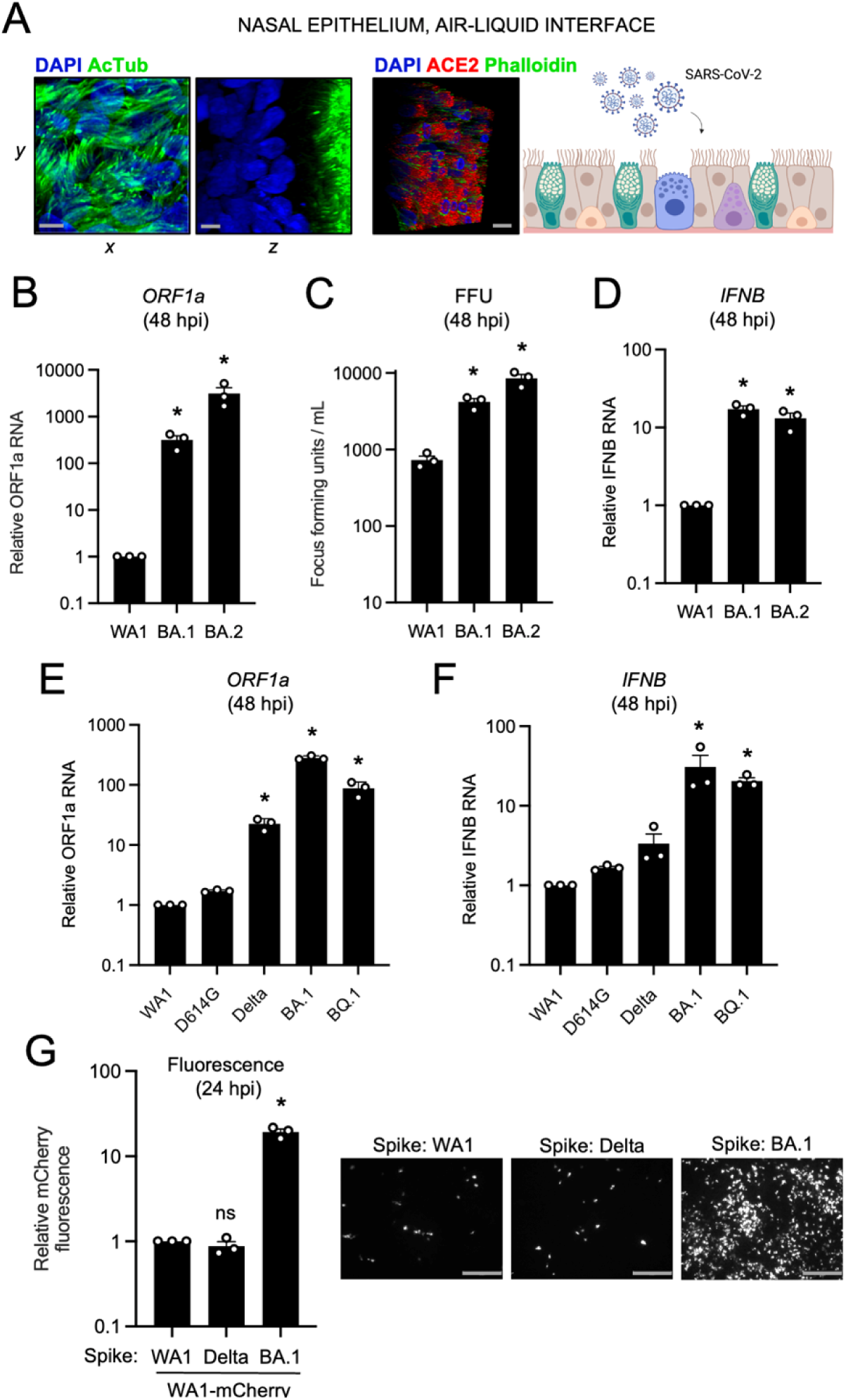
Omicron Spike enables enhanced SARS-CoV-2 infectivity in primary nasal epithelia cultured at the air-liquid interface. (A) Primary human nasal epithelial cells (pooled from 14 human donors) were cultured at the air-liquid interface to mimic the human respiratory tract and challenged with SARS-CoV-2. Acetylated tubulin levels were determined by anti-AcTub immunofluorescence, ACE2 levels were determined by anti-ACE2 immunofluorescence, and DAPI was used to stain nuclei. 3D reconstructions of *xy* and *yz* fields are shown. Scale bars = 10 µm. Cartoon made with Biorender.com. (B) Cells were subjected to total cellular RNA extraction at 48 hours post inoculation with 10000 plaque forming units of WA1, BA.1, or BA.2, and RT-qPCR of viral ORF1a was performed. Relative ORF1a transcript abundance compared to actin was determined by the 2^(-ΔΔCT) method. ORF1a abundance of WA1 at 48 hours post inoculation was set to 1. (C) Infectious virus titers from cell culture medium collected at 48 hours post-inoculation were measured by challenging Vero E6-TMPRSS2 cells followed by fixation and immunostaining with anti-N antibody. Infectious units were quantified by measuring focus forming units with high-content imaging. Symbols represent results from three independent infections of Vero E6-TMPRSS2 cells. (D) RT-qPCR of cellular *IFNB* was performed at 48 hours post inoculation. Relative *IFNB* transcript abundance compared to actin was determined by the 2^(-ΔΔCT) method. *IFNB* abundance of WA1 at 48 hours post inoculation was set to 1. (E) Viral ORF1a RT-qPCR was performed at 48 hours post-inoculation with 10000 plaque forming units of WA1, D614G, Delta, BA.1, or BQ.1. Relative ORF1a transcript abundance compared to actin was determined by the 2^(-ΔΔCT) method. ORF1a abundance of WA1 at 48 hours post inoculation was set to 1. (F) RT-qPCR of cellular *IFNB* was performed at 48 hours post inoculation. Relative *IFNB* transcript abundance compared to actin was determined by the 2^(-ΔΔCT) method. *IFNB* abundance of WA1 at 48 hours post inoculation was set to 1. (G) Recombinant WA.1 encoding mCherry and Spike protein from WA1, Delta, or BA.1 (WA1-mCherry (WA1 Spike), WA1-mCherry (Delta Spike), and WA1-mCherry (BA.1 Spike)) were used to inoculate primary human nasal epithelial cells (pooled from 14 human donors) cultured at the air-liquid interface, and infection was measured by mCherry fluorescence at 24 hours post-inoculation. Summary mCherry fluorescence from three independent infections is shown. The fluorescence intensity of WA1 (WA1 Spike)-mCherry was set to 1. Representative fields of view are shown. Scale bars = 300 µm. All results are represented as means plus standard error from three independent infections (symbols represent biological replicates). Statistically significant differences (* *P* < 0.05) between the indicated condition and the corresponding data point of WA.1 were determined by one-way ANOVA.

Given the degree to which Omicron BA.1 and BA.2 replicate in primary nasal ALI, we also tested the infective potential of an Omicron sublineage more recently circulating in humans (BQ.1) as well as Delta and the D614G variant, the ancestor of Pango lineage B that gave rise to all SARS-CoV-2 variants of concern. D614G replicated to a similar extent as WA1, while Delta achieved a significantly higher degree of replication (23-fold). However, replication of Omicron lineages BA.1 and BQ.1 in nasal ALI far exceeded that of all variants tested (282- and 88-fold, respectively) (**Figure 2E**). While all viruses triggered the expression of the interferon-stimulated gene *IFITM3* (**Supplemental Figure 2B**), BA.1 and BQ.1 induced more *IFNB* expression compared to the other variants (**Figure 2F**). We also inoculated nasal ALI with Omicron XBB, the ancestor of sublineages currently circulating in humans as of October 2023, and it replicated to a significantly greater extent than D614G (**Supplemental Figure 2C**).

To determine whether the enhanced nasal tropism of Omicron BA.1 can be attributed to its Spike protein, we generated mCherry-expressing, recombinant SARS-CoV-2 WA1 encoding Spike from WA1, Delta, or BA.1 (referred to as WA1 (WA.1 Spike), WA1 (Delta Spike), or WA1 (BA.1 Spike). We found that the presence of Spike from BA.1, but not Delta, resulted in a significant gain in nasal cell infectivity relative to WA1 (20-fold by mCherry fluorescence (**Figure 2G**) or 100-fold by viral ORF1a RT-qPCR at 24 hours post-inoculation) **(Supplemental Figure 2D)**. Results were similar regardless of how we calculated virus titers to prepare inoculants (infection of Vero E6-TMPRSS2 cells (**Supplemental Figure 2D**) or viral ORF1a RT-qPCR (**Supplemental Figure 2E**). When earlier timepoints were examined, WA1 (BA.1 Spike)-infected cells were observed as early as 6 hours post-inoculation and numbers increased substantially between 9 and 12 hours post-inoculation, indicative of rapid virus spread in nasal ALI (**Supplemental Figure 2F**). These results formally indicate that the elevated nasal tissue infectivity of BA.1 is governed by its Spike protein, and more specifically, by the mutations that distinguish it from Delta Spike. Therefore, enhanced nasal cell tropism evolved recently in the Omicron ancestor as a result of unique mutations in Spike.

### Entry of Omicron BA.1 into primary nasal epithelial cells is mediated by proteases of the MMP/ADAM families and occurs independently of endocytosis

Since BA.1 Spike encodes for an unprecedented ability to enter human nasal epithelia, we characterized the subcellular location at which Omicron Spike-mediated entry occurs in nasal epithelia and determined the cellular proteases that enable it. Several reports claim that Omicron enters cells via cathepsin-dependent fusion in endosomes^31,32,36^, but the Omicron entry pathway may vary in transformed cell lines^44^. Therefore, we examined the entry determinants of WA1, Delta, and BA.1 in primary human nasal ALI. We used E64d to disrupt cathepsin-mediated entry in endolysosomes and found that, while WA1 replication was reduced by 25%, that of Delta and BA.1 were not **(Figure 3A)**. Interestingly, Delta and BA.1 infections were enhanced approximately 2-fold by E64d treatment, despite the same dose being previously shown to partially inhibit Omicron infection in a transformed lung cell line^31^. However, addition of camostat mesylate (an inhibitor of serine transmembrane proteases including TMPRSS2) alone or in combination with E64d strongly blocked infection of WA1 and Delta in nasal ALI **(Figure 3A)**. These data suggest that WA1 and Delta enter nasal ALI primarily through a TMPRSS2-dependent entry route at the plasma membrane. In contrast, infection of nasal ALI by BA.1 was completely insensitive to blockade by camostat **(Figure 3A)**, suggesting the use of a divergent entry pathway that is TMPRSS2-independent. The TMPRSS2-independence of Omicron infection has been reported by multiple groups, and as a result, it has been inferred that Omicron uses an endosomal, cathepsin-dependent route to enter human cells^31,32,36^. However, a recently described metalloproteinase-mediated entry pathway for SARS-CoV-2 that may enable virus fusion at the plasma membrane^11,45^ prompted us to test the effect of incyclinide, a broad-spectrum inhibitor of MMP and ADAM metalloproteinases. WA1, Delta, and BA.1 infections were reduced by incyclinide treatment in primary ALI, but BA.1 was particularly sensitive (inhibited by 27-fold compared to 4-fold inhibition of WA1 and 5-fold inhibition of Delta) **(Figure 3A)**. These findings suggest that processing of BA.1 Spike by metalloproteinases, but not by TMPRSS2, enables BA.1 entry into primary nasal epithelia. To address whether a metalloproteinase-dependent route of entry allows for fusion at the plasma membrane or at endolysosomal membranes, we blocked endosomal trafficking with the endolysosomal acidification inhibitor bafilomycin A1. WA1 and Delta infections were inhibited by bafilomycin A1, suggesting that these viruses can utilize an endocytic pathway for entry (**Figure 3A**). In contrast, bafilomycin A1 enhanced BA.1 infection 4-fold, suggesting that BA.1 utilizes an entry pathway that is non-endocytic **(Figure 3A)**. We also tested the inhibitor sensitivity of recombinant WA1 (Delta Spike) and WA1 (BA.1 Spike) to further clarify the entry pathway unlocked by BA.1 Spike in nasal ALI. As seen with the full-length clinical isolates, WA1 (Delta Spike) was inhibited by bafilomycin A1 while WA1 (BA.1 Spike) was not (**Figure 3B**). However, WA1 (BA.1 Spike) was inhibited 10-fold by combined treatment of bafilomycin A1 and incyclinide. For WA1 (Delta Spike), treatment with bafilomycin A1 and incyclinide inhibited infection to the same extent as bafilomycin A1 alone (**Figure 3B**). These results may suggest that metalloproteinases at the cell surface promote entry mediated by BA.1 Spike, while those in endosomes may promote entry mediated by Delta Spike. Overall, our data support a model whereby Delta Spike may be processed by TMPRSS2 at the plasma membrane or by metalloproteinases in endosomes, allowing for fusion at both sites in nasal ALI. In contrast, BA.1 Spike may be processed primarily by metalloproteinases at or near the cell surface and fusion ensues in this compartment, not in endosomes.

**Figure 3.**
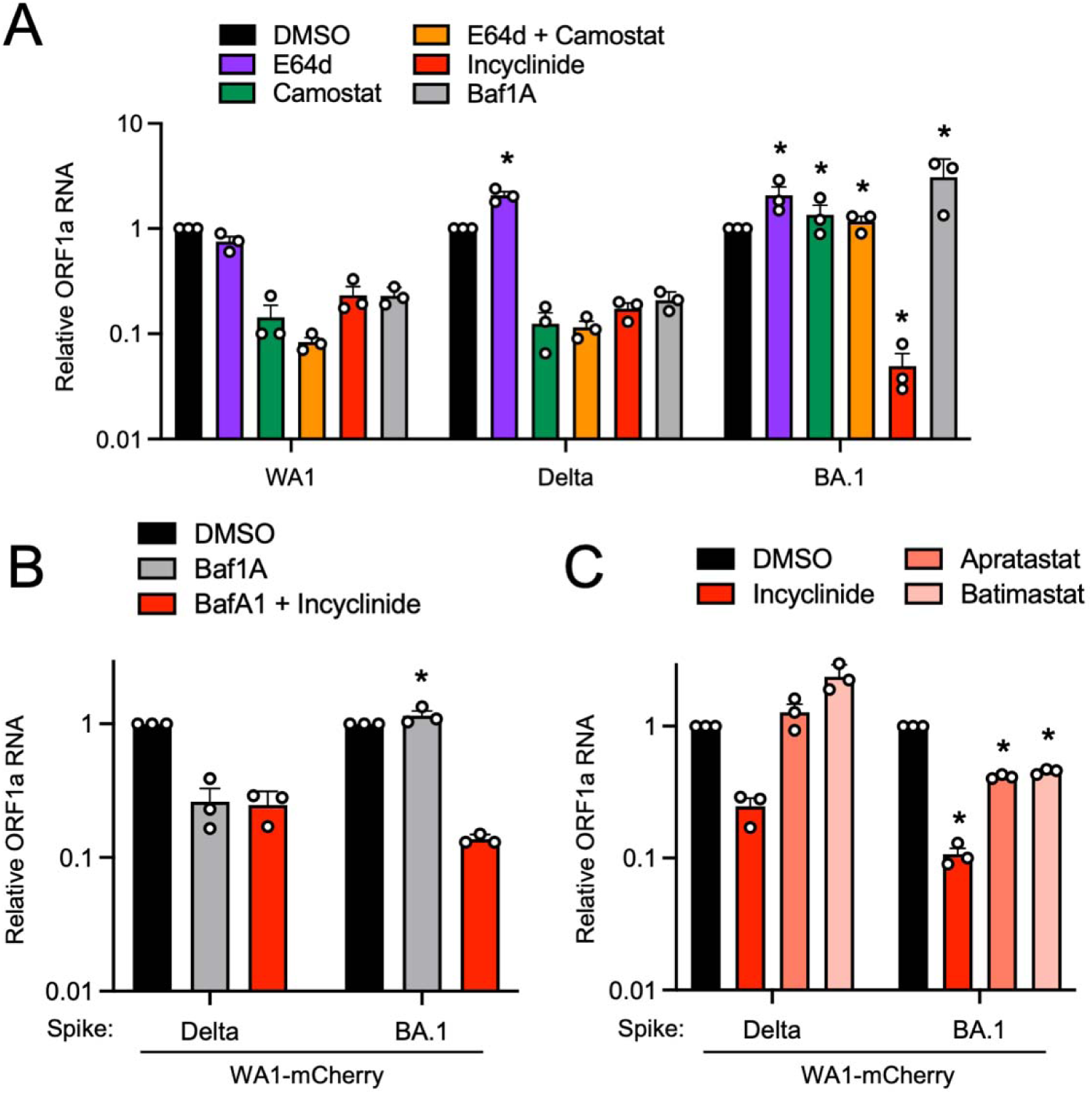
Entry of Omicron BA.1 into primary nasal epithelial cells is mediated by proteases of the MMP/ADAM families and occurs independently of endocytosis. (A) Primary human nasal epithelial cells (pooled from 14 human donors) were cultured at the air-liquid interface and pretreated with 10 µM E64d, 10 µM camostat mesylate, a combination of 10 µM E64d and 10 µM camostat mesylate, 10 µM incyclinide, or 1 µM bafilomycin A1 for 2 hours. Afterwards, cells were challenged with WA.1, Delta, or BA.1 at an MOI of 0.05. At 48 hours post inoculation, cells were subjected to total cellular RNA extraction and ORF1a levels were measured by RT-qPCR. Relative ORF1a transcript abundance compared to actin was determined by the 2^(-ΔΔCT) method. For each virus, ORF1a abundance in the DMSO condition was set to 1. All results are represented as means plus standard error from three independent infections (symbols represent biological replicates). Statistically significant differences (* *P* < 0.05) between the indicated condition and the corresponding condition of WA.1 were determined by one-way ANOVA. ORF1a abundance in the DMSO condition at 48 hours post inoculation was set to 1. (B) Primary human nasal epithelial cells (pooled from 14 human donors) were cultured at the air-liquid interface and pretreated with 1 µM bafilomycin A1 or 1 µM bafilomycin A1 and 10 µM incyclinide for 2 hours. Afterwards, cells were challenged with WA1-based recombinant viruses encoding Delta Spike or BA.1 Spike at an MOI of 0.05. At 48 hours post-inoculation, ORF1a levels were measured by RT-qPCR. For each virus, ORF1a abundance in the DMSO condition was set to 1. (C) Primary human nasal epithelial cells (pooled from 14 human donors) were cultured at the air-liquid interface and pretreated with 10 µM incyclinide, 10 µM aprastatat, or 10 µM batimastat for 2 hours. Afterwards, cells were challenged with WA1-based recombinant viruses encoding Delta Spike or BA.1 Spike at an MOI of 0.05. At 48 hours post-inoculation, ORF1a levels were measured by RT-qPCR. For each virus, ORF1a abundance in the DMSO condition was set to 1. Statistically significant differences (* *P* < 0.05) between the indicated condition of BA.1 and the corresponding condition of Delta were determined by one-way ANOVA. All results are represented as means plus standard error from three independent infections (symbols represent biological replicates).

To further substantiate a role for metalloproteinases in the entry process driven by different Spikes in nasal ALI, we tested the sensitivity of WA (Delta Spike) and WA1 (BA.1 Spike) to additional metalloproteinase inhibitors. Apratastat is an inhibitor of ADAM10, ADAM17 and MMP-13, and batimastat is an inhibitor of MMP-1, MMP-2, MMP-3, MMP-7, MMP-9, ADAM10, and ADAM17. While incyclinide inhibited WA1 (Delta Spike) infection, apratastat and batimastat did not inhibit but instead modestly boosted infection (**Figure 3C**). In contrast, both apratastat and batimastat reduced WA (BA.1 Spike) infection by 60% by (**Figure 3C**). These results reveal a functional divergence between Delta Spike and BA.1 Spike in terms of their processing by metalloproteinases. Our results suggest that, at the concentrations used, apratastat and batimastat inhibit a narrower range of metalloproteinases than incyclinide. Nonetheless, the elevated sensitivity of WA1 (BA.1 Spike) to inhibition by apratastat and batimastat suggests that BA.1 Spike can be processed by metalloproteinases targeted by both compounds—such as ADAM10 and ADAM17. Furthermore, since apratastat and batimastat decrease BA.1 Spike-mediated infection, ADAM10 and ADAM17 may be elements of a productive entry pathway for BA.1 in nasal ALI. In contrast, processing of Delta Spike by these metalloproteinases appears to have deleterious consequences for Delta entry, since apratastat and batimastat boosted infection rather than inhibited it (**Figure 3C**). Therefore, BA.1 Spike has evolved to utilize a unique entry route that is, while TMPRSS2-independent, dependent upon ADAM10, ADAM17, and other metalloproteinases in the nasal epithelia. To assess whether preferential usage of neuropilin-1 at the cell surface by BA.1 Spike may explain the increased adherence of BA.1 to nasal cell epithelia and adoption of a unique entry pathway, we tested the impact of the neuropilin-1 inhibitor EG00229 on nasal ALI infection by WA1 (WA1 Spike), WA1 (Delta Spike), and WA1 (BA.1 Spike). WA1-Spike was most sensitive to inhibition by EG00229, while Delta Spike was completely resistant, and BA.1 Spike conferred modest sensitivity (**Supplemental Figure 3**). These results indicate that the extent of neuropilin-1 dependency does not explain the differential ability for WA1 Spike, Delta Spike, and BA.1 Spike to mediate entry into nasal epithelia. Therefore, factors other than neuropilin-1 may be responsible for the improved infectivity of BA.1 in nasal tissue.

### Omicron Spike enables an entry pathway into primary nasal epithelial cells that is resistant to inhibition by type-I and type-III interferons

Next, we explored whether the increased nasal cell infectivity exhibited by Omicron is related to evasion of or resistance to cell-intrinsic antiviral defenses. During the preparation of this manuscript, it was demonstrated that SARS-CoV-2 variants of concern displayed an increasing resistance to inhibition by interferon (IFN) treatment, with Omicron exhibiting the greatest degree of resistance ^46^. However, the role of Spike in conferring IFN resistance was not assessed, and it was unknown whether an Omicron-specific cellular entry pathway enabled IFN-resistance in nasal epithelia. Therefore, we tested whether the unique entry requirements of Omicron Spike allow for evasion of the IFN-induced antiviral state in primary nasal ALI (**Figure 4A**). Pre-treatment with IFN-beta or IFN-lambda inhibited WA1 in a dose-dependent manner (up to two orders of magnitude) as measured by viral ORF1a RT-qPCR, while BA.1 or BA.2 infections were significantly less impacted **(Figure 4B)**. We also quantified infectious virus yield under these conditions and confirmed that BA.1 and BA.2 achieved higher titers and were inhibited by IFN to a significantly lesser extent than WA1 **(Figure 4C)**. When we compared the sensitivity of full-length clinical virus isolates with recombinant WA1 encoding different Spike proteins, we found that the identity of Spike governed sensitivity to IFN. Compared to WA1, which was inhibited 6-fold by low amounts of IFN-beta or IFN-lambda, WA1 (Delta Spike) exhibited a slight decrease in sensitivity to IFN (**Figure 4D**). However, WA1 (BA.1 Spike) was completely insensitive to the same doses of IFN, and this closely resembled the resistance profile of full-length BA.1 (**Figure 4D**). Therefore, BA.1 Spike is sufficient to confer IFN-resistance to an otherwise IFN-sensitive virus, while Delta Spike is not. Moreover, these results reveal the presence of host factors intrinsic to nasal cells that are interferon-inducible and target the Spike-mediated entry step.

**Figure 4.**
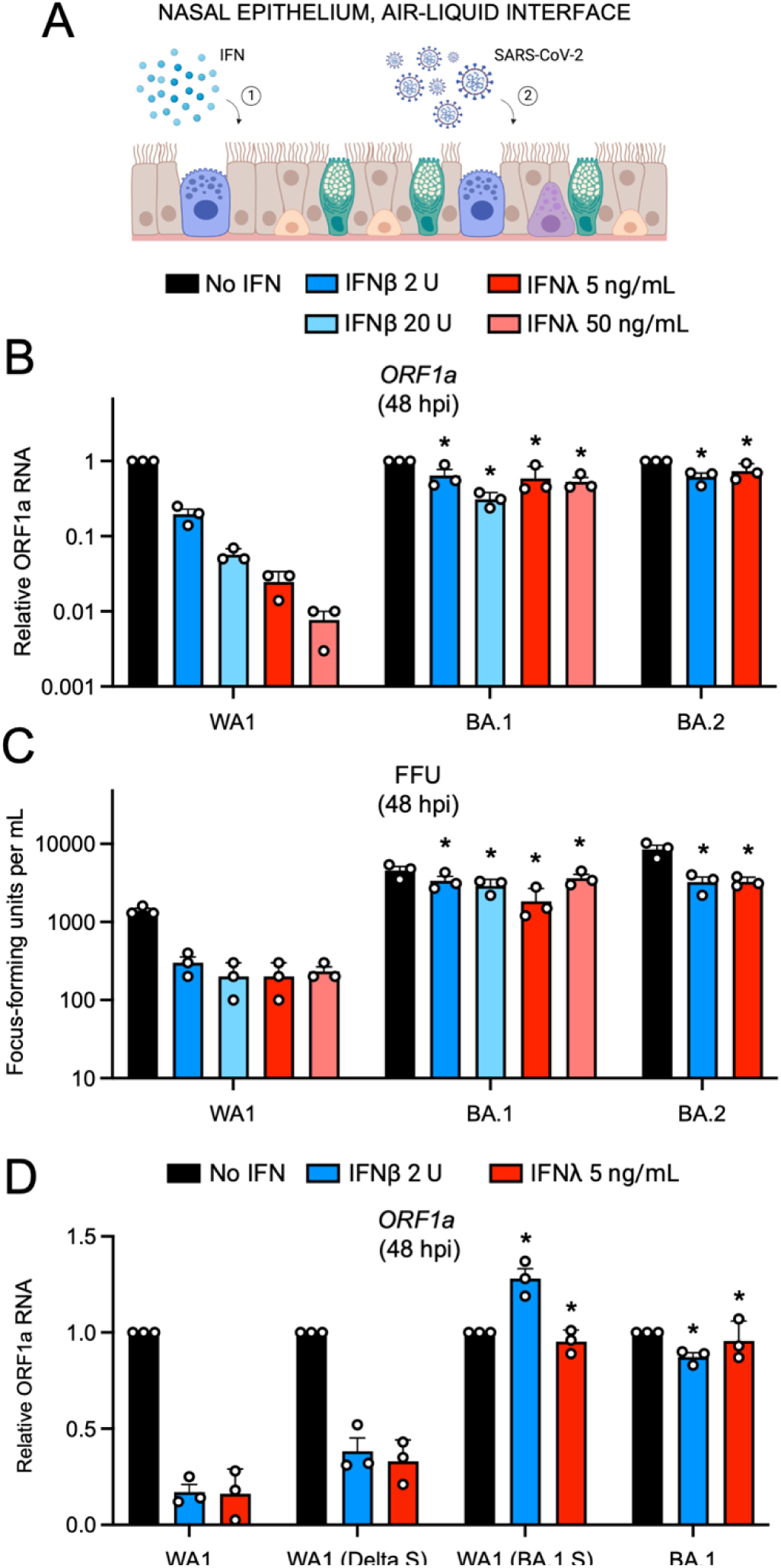
Omicron Spike enables an entry pathway into primary nasal epithelial cells that is resistant to inhibition by type-I and type-III interferons. (A) Primary human nasal epithelial cells (pooled from 14 human donors) were cultured at the air-liquid interface, pre-treated for 18 hours with the indicated amounts of IFN-beta or IFN-lambda, and challenged with 10000 plaque forming units of WA1, BA.1, or BA.2. Cartoon made with Biorender.com. (B) At 48 hours post inoculation, total RNA was extracted from cells and subjected to RT-qPCR. ORF1a levels were measured compared to actin by the 2^(-ΔΔCT) method. For each virus, ORF1a abundance in the No IFN condition was set to 1. (C) At 48 hours post inoculation, infectious virus titers from cell culture medium were measured by challenging Vero E6-TMPRSS2 cells followed by fixation and immunostaining with anti-N antibody. Infectious units were quantified by measuring focus forming units by high-content imaging. Symbols represent independent infections of Vero E6-TMPRSS2 cells. (D) Primary human nasal epithelial cells (pooled from 14 human donors) were cultured at the air-liquid interface, pre-treated for 18 hours with the indicated amounts of IFN-beta or IFN-lambda, and challenged with 10000 plaque forming units of WA1, WA1-mCherry (Delta Spike), WA1-mCherry (BA.1 Spike), or BA.1. ORF1a levels were measured by RT-qPCR and compared to actin by the 2^(-ΔΔCT) method. For each virus, ORF1a abundance in the No IFN condition was set to 1. All results are represented as means plus standard error from three independent infections (symbols represent biological replicates). Statistically significant differences (* *P* < 0.05) between the indicated condition and the corresponding data point of WA.1 were determined by one-way ANOVA.

We also examined the IFN sensitivity of WA1, BA.1 and BA.2 in submerged primary nasal monolayers **(Supplemental Figure 4A)**. While IFN-beta or IFN-lambda exposure strongly blocked WA1 infection by more than 100-fold, the same amount of IFN inhibited BA.1 and BA.2 to a much lesser extent (less than 2-fold) **(Supplemental Figure 4B)**. To confirm that Spike is a major determinant of IFN sensitivity in these cells, we generated pseudovirus consisting of vesicular stomatitis virus (VSV) decorated with Spike from WA1, BA.1, or BA.2. VSV-WA1 Spike was exquisitely sensitive to inhibition by IFN-beta or IFN-lambda in a dose-dependent manner, with inhibition up to two orders of magnitude observed (**Supplemental Figure 4C**). In contrast, VSV-BA.1 Spike and VSV-BA.2 Spike were much less sensitive to inhibition by IFN-beta or IFN-lambda **(Supplemental Figure 4C)**. Therefore, Spike is the major determinant of the IFN resistance displayed by Omicron sublineages, and this feature can be transferred to other viruses by pseudotyping. Additionally, we determined that cellular attachment of WA1 and BA.1 in nasal monolayers was not affected by IFN-beta or IFN-lambda, indicating that nasal cell-intrinsic host defenses act at a post-attachment step of the virus entry process **(Supplemental Figure 4D)**. Therefore, Omicron Spike enables evasion of restriction by nasal cell-intrinsic host factors inhibiting the post-attachment step of virus entry.

### Amphotericin B partially relieves a block to infection mediated by WA1 Spike and Delta Spike in primary nasal epithelia

A limited number of interferon-stimulated genes are known to block the cellular entry step of enveloped viruses^47^. Among the best characterized are the interferon-inducible transmembrane (IFITM) proteins. IFITM proteins provide constitutive and interferon-induced antiviral protection in many cell types by inhibiting the virus-cell membrane fusion step required for entry^47,48^. While it is established that murine IFITM3 restricts SARS-CoV-2 infection and limits viral disease in vivo^49,50^, the roles played by human IFITM proteins during SARS-CoV-2 infection have remained controversial. For example, it was demonstrated that IFITM proteins can inhibit SARS-CoV-2 infection in endosomes but can promote infection at the cell surface^51,52^. Furthermore, it has been suggested that endogenous human IFITM proteins in lung epithelial cells enable SARS-CoV-2 infection by possibly acting as a receptor or attachment factor^53^. To address whether IFITM proteins act as a barrier to SARS-CoV-2 infection in primary human nasal epithelia, we tested the impact of the antifungal Amphotericin B on infection with recombinant WA1 (WA1 Spike), WA1 (Delta Spike), and WA1 (BA.1 Spike). Amphotericin B was previously reported to counteract the antiviral properties of human IFITM2 and IFITM3^54-56^ by restoring membrane fluidity in cellular membranes where IFITM3 is present^57^. In nasal ALI pretreated with Amphotericin B or left untreated, we inoculated with WA1 (WA1 Spike), WA1 (Delta Spike), or WA1 (BA.1 Spike) and scored infection by mCherry fluorescence at 24 hours post-inoculation. As described above, WA1 (BA.1 Spike) displayed a far superior capacity to infect untreated nasal ALI relative to WA1 (WA1 Spike) and WA1 (Delta Spike) (**Figure 5**). However, Amphotericin B treatment led to marked increases in WA1 (WA1 Spike) and WA1 (Delta Spike) infections, boosting them by approximately 7-fold. By comparison, Amphotericin B had a negligible impact on WA1 (BA.1 Spike) infection at the same time point (**Figure 5**). We also determined whether Amphotericin B impacted WA1 (BA.1 Spike) infection at earlier time points when infected cells were less abundant. At 9 and 12 hours post-inoculation, Amphotericin B marginally boosted infection of WA1 (BA.1 Spike) (less than 2-fold) (**Supplemental Figure 5**). Therefore, these results suggest that human IFITM proteins restrict WA1 (WA1 Spike) and WA1 (Delta Spike) in primary nasal epithelia, while BA.1 Spike confers escape from this restriction. Therefore, antiviral IFITM proteins form a part of the nasal cell-intrinsic defenses that limit infection by early/ancestral SARS-CoV-2.

**Figure 5.**
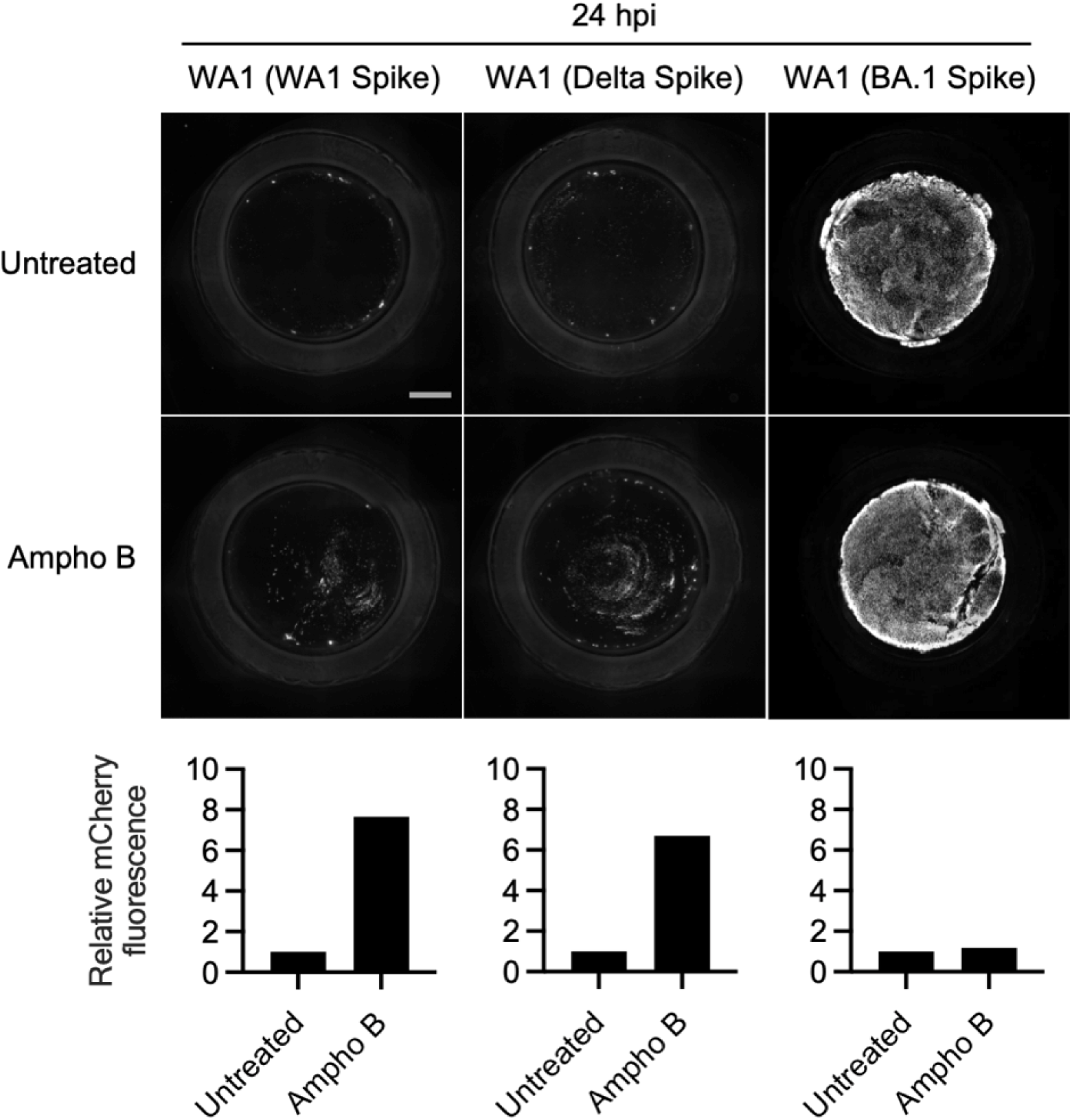
Amphotericin B partially relieves a block to infection mediated by WA1 Spike and Delta Spike in primary nasal epithelia. Primary human nasal epithelial cells (pooled from 14 human donors) were cultured at the air-liquid interface, pre-treated with 1 µM Amphotericin B for two hours or untreated, and inoculated with 10000 plaque forming units of WA1-mCherry (WA1 Spike), WA1-mCherry (Delta Spike), or WA1-mCherry (BA.1 Spike). mCherry fluorescence was measured at 24 hours post-inoculation by high-content imaging. The fluorescence intensity of the untreated condition for each virus was set to 1. Scale bar = 300 µm. Ampho B; amphotericin B.

Overall, by combining experiments with authentic SARS-CoV-2, recombinant virus, and Spike-decorated pseudovirus, we show that Spike is the major determinant of the enhanced infectivity and interferon resistance exhibited by Omicron in primary human nasal epithelia, two phenotypes which are associated with a unique dependence on metalloproteinases for cell entry at or near the cell surface.

## Discussion

Using entirely primary nasal epithelia from human adult donors, we demonstrate that Omicron (including BA.1, BA.2, BQ.1, and XBB) exhibits drastically increased infectivity in nasal tissue compared to early SARS-CoV-2, the D614G variant, and the Delta variant of concern. This mapped to Omicron Spike and, at least in part, to an increased ability for Omicron virions to adhere to the surface of nasal epithelia, which occurred independently of the presence of cilia. Furthermore, we show that Omicron utilizes an entry route into cells that depends on cellular metalloproteinases but that is independent of the transmembrane serine protease TMPRSS2. Since we also demonstrate that Omicron Spike-mediated infection enables evasion of the antiviral state induced by type-I and type-III IFN in this tissue, these results suggest that the metalloproteinase-dependent entry pathway utilized by Omicron may promote escape from constitutive and IFN-induced antiviral factors targeting virus entry.

During the preparation of this manuscript, it was reported that BA.1 exhibits a growth advantage in primary human nasal epithelia compared to the Delta variant^31^. Here, using recombinant SARS-CoV-2, we extend this finding by demonstrating that BA.1 Spike is responsible for increased nasal cell infectivity. However, in another study, while BA.1 exhibited a gain in nasal cell infectivity compared to Alpha, no significant differences in nasal cell infectivity were observed between Delta and BA.1 ^32^. This discrepancy is likely the result of utilization of distinct cell culture models to compare relative infectivity. Our use of recombinant, fluorogenic SARS-CoV-2 in nasal ALI derived from pools of 14 human adult donors shows that Spike from BA.1 promotes a gain in nasal cell infectivity compared to Delta Spike. Therefore, the emergence and evolution of Omicron involved unique mutations in Spike that contributed to enhanced nasal cell binding and entry, and this may explain how Omicron replaced previously circulating variants of concern and continues to persist in human populations.

It was previously reported that human nasal airways express relatively low levels of TMPRSS2, and combined with the demonstration that Omicron exhibits TMPRSS2-independence in transformed cell lines, others have inferred that Omicron is more reliant on endosomal cathepsins for entry ^32^. However, our findings with primary nasal epithelia reveal that TMPRSS2-independence does not necessarily imply the use of an endosomal entry route by Omicron. Metalloproteinases such as MMP-14, MMP-16, ADAM10, and ADAM17 are also highly expressed in upper airways and have been shown to activate Omicron Spike-mediated fusion at the surface of several transformed cell lines ^11^. Here, we demonstrate that the primary route of entry of Omicron in primary nasal epithelia is mediated by metalloproteinases. The pan-metalloproteinase inhibitor incyclinide strongly blocked BA.1 infection in nasal ALI, but the endosomal cathepsin inhibitor E64d and the endolysosomal acidification inhibitor bafilomycin A1 did not. Instead, these endosomal inhibitors boosted infection of BA.1 in nasal epithelia, suggesting that the endosomal entry route is deleterious for BA.1 in this tissue. Bafilomycin A1 inhibits vacuolar-ATPase to prevent endolysosomal acidification, and furthermore, endosomal maturation and transport between early and late endosomes is repressed ^58^. Therefore, our results suggest that processing of Spike by metalloproteinases enables fusion at the plasma membrane and/or in early or recycling endosomes, while late endosomes or lysosomes and the cathepsins found therein are unlikely to be involved.

One possibility explaining the enhanced capacity for Omicron Spike to adhere to and enter nasal cells (either in the presence or absence of ciliated cells) is its increase in net overall positive charge compared to previous variants of concern^59^. It has also been proposed that Omicron Spike may adhere more strongly to attachment factors present at the epithelial cell surface, including heparan sulfate^60,61^ and neuropilin-1^62^. In this study, sensitivity to neuropilin-1 inhibitor did not correlate with the nasal cell infectivity of WA1, Delta, or BA.1, suggesting that other cellular factors may play a role (**Supplemental Figure 3**). A significant aim of future research will be to establish whether mutations in Omicron Spike promote increased residence time at the cell surface and whether this influences its dependence on certain cellular proteases for cleavage and the triggering of fusion. It is possible that longer residency at the cell surface may negatively interfere with processing by TMPRSS2 and steer processing towards metalloproteinases.

Another factor that likely contributed to the spread and dominance of Omicron lineages is their resistance to the antiviral properties of IFN. Early/ancestral SARS-CoV-2 was previously reported to antagonize IFN signaling in cells, and nearly every viral protein encoded by SARS-CoV-2 has been shown to contribute to evasion of IFN through various mechanisms^63-65^. While the relative insensitivity of Omicron to exogenous administration of multiple IFN types has been reported previously^46,66^, the viral proteins responsible for rendering IFN ineffective against Omicron were unknown. Other studies showed that Omicron exhibits a decreased capacity to interfere with IFN production and signaling, relative to early/ancestral SARS-CoV-2 or Delta^44,67-69^, and our measurements of type-I IFN production by Omicron-infected nasal ALI affirm this finding. However, in one case, Omicron was purported to induce a relatively inferior IFN response compared to Delta ^66^, and this discrepancy underlines the importance of assessing this activity in primary human upper airway epithelia. Overall, we postulate that Omicron exhibits resistance to IFN not because of its ability to interfere with the production or function of IFN from infected cells, but rather, because Omicron Spike enables passage into nasal cells in a manner which is insensitive to IFN-stimulated proteins. In addition, it is possible that Omicron displays enhanced infectivity in nasal cells even in the absence of IFN because Omicron Spike confers resistance to antiviral factors that are constitutively expressed in nasal epithelia and further upregulated by type-I or type-III IFN. Our results indicating that Amphotericin B partially relieves a constitutive barrier to infection mediated by IFITM proteins provides direct evidence for this possibility. Furthermore, our findings provide a new explanation for why Omicron exhibits decreased IFN antagonism—if the Spike-mediated entry process of Omicron is resistant to type-I and type-III IFN, the selective advantage to antagonize IFN pathways may be lost. Furthermore, the improved capacity for Omicron lineages to infect nasal cells despite the induction of an antiviral state therein provides a plausible scenario for how Omicron displaced previous variants of concern. It was demonstrated that IFN produced from Omicron-infected cells can inhibit infection by Influenza A virus ^67^, so it is plausible that the IFN response elicited by Omicron infection may render nasal epithelia resistant to infection by previous SARS-CoV-2 variants of concern.

Our description of the IFN resistance conferred by Omicron Spike has implications for the ongoing development of type-I and type-III IFN as antiviral therapeutics for the prevention or treatment of SARS-CoV-2 infection and COVID-19. A case has been made for the deployment of type-III IFN in the context of SARS-CoV-2 infection, since the receptor for type-III IFN is expressed in the respiratory tract but is not widely distributed elsewhere (resulting in fewer unintended side effects such as inflammation)^63,70^. As an important proof-of-concept, nasal administration of IFN-lambda was protective against SARS-CoV-2 variants, including Omicron, in mice ^71^. However, clinical trials thus far in humans have reported conflicting findings. One such study reported that participants who were administered a single injection of IFN-lambda following symptom onset showed no significant reductions in viral shedding or symptom severity compared to placebo^72^, while another reported that a single injection upon symptom onset resulted in reduced viral loads^73^ and reduced COVID-19-related hospitalizations^74^. However, in neither case was it determined how IFN-lambda fared in individuals infected with Omicron variants. Since Omicron displays a greater degree of IFN-resistance compared to earlier variants, additional clinical studies are needed to test the suitability of IFN-lambda in the fight against contemporaneous Omicron and additional variants that arise in the future.

While some mutations in Omicron Spike may have conferred Omicron with superior evasion of vaccine-elicited neutralizing antibodies compared to the Delta variant, our findings indicate that mutations in Omicron Spike also allow for efficient seeding of infection in the nasal epithelium and resistance to the interferon-induced antiviral state therein. Our findings provide mechanistic insight into the basis for the efficient transmission and persistence of Omicron in vaccinated and unvaccinated human populations. Future efforts aimed at understanding how individual Spike mutations contribute to these different phenotypes may enable a better understanding of the selective forces driving their emergence.

## Figure Legends

**Supplemental Figure 1.**
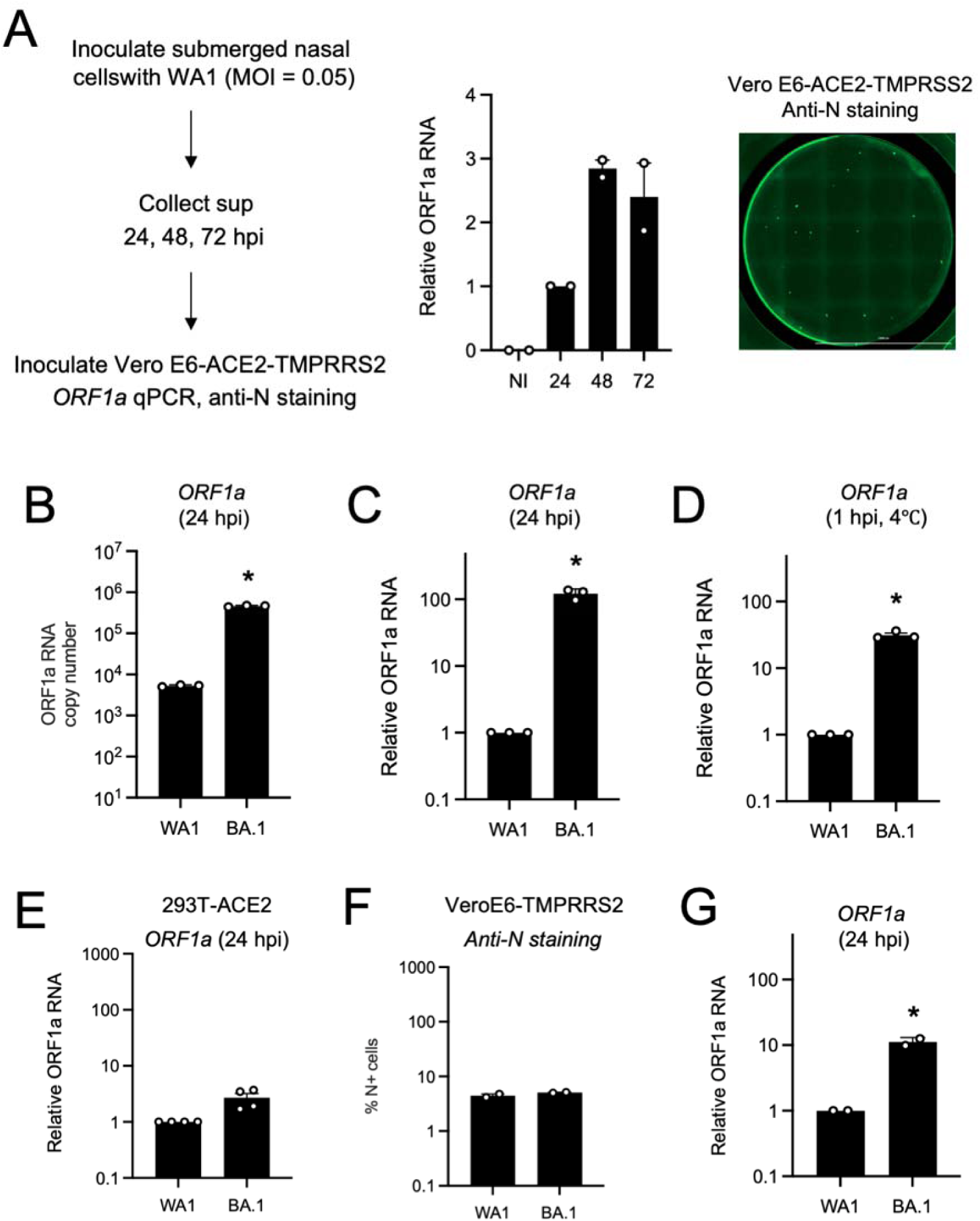
(A) Primary human nasal epithelial cells (pooled from 3 human donors) were cultured as undifferentiated, submerged monolayers and challenged with WA1 at an MOI of 0.05. Cell culture supernatants were collected at 24, 48, and 72 hours post-inoculation and added to Vero E6-ACE2-TMPRSS2 cells. 24 hours later, Vero E6-ACE2-TMPRSS2 cells were subjected to total RNA extraction and viral ORF1a RT-qPCR was performed. In addition, productive infection of Vero E6-ACE2-TMPRSS2 by the 24 hours post-inoculation supernatant was confirmed by anti-N immunofluorescence microscopy. (B) Primary human nasal epithelial cells (cells from three human donors, pooled) were inoculated with WA1 or BA.1 at an MOI of 0.05. Total cellular RNA was extracted and viral ORF1a was quantified by RT-qPCR at 24 hours post-inoculation. Absolute ORF1a RNA copy numbers were calculated by comparison to an ORF1a standard curve. (C) Relative viral ORF1a RNA abundance compared to actin was determined by the 2^(-ΔΔCT) method. ORF1a abundance of WA1 was set to 1. (D) Primary human nasal epithelial cells (cells from three human donors, pooled) were inoculated with WA1 or BA.1 at an MOI of 0.05 on ice. At 1 hour post-inoculation, total cellular RNA was extracted and viral ORF1a was quantified by RT-qPCR to measure virus adherence to cells. Relative viral ORF1a RNA abundance compared to actin was determined by the 2^(-ΔΔCT) method. ORF1a abundance of WA1 was set to 1. (E) HEK293T-ACE2 cells were inoculated with WA1 or BA.1 at an MOI of 0.05. 24 hours post-inoculation, total cellular RNA was extracted and viral ORF1a was quantified by RT-qPCR. Relative viral ORF1a RNA abundance compared to actin was determined by the 2^(-ΔΔCT) method. ORF1a abundance of WA1 was set to 1. (F) Vero E6-TMPRSS2 cells were inoculated with WA1 or BA.1 at an MOI of 0.05. 24 hours post-inoculation, cells were fixed, stained with anti-N antibody, and infection was scored by flow cytometry. (G) Primary human nasal epithelial cells (cells from three human donors, pooled) were inoculated with WA1 or BA.1 (5×10^7 copies of absolute ORF1a RNA used as input). 24 hours post-inoculation, total cellular RNA was extracted and viral ORF1a was quantified by RT-qPCR. Relative viral ORF1a RNA abundance compared to actin was determined by the 2^(-ΔΔCT) method. ORF1a abundance of WA1 was set to 1. All results are represented as means plus standard error from three independent infections (symbols represent biological replicates). Statistically significant differences (* *P* < 0.05) between the indicated condition of BA.1 and the corresponding condition of WA.1 were determined by one-way ANOVA.

**Supplemental Figure 2.**
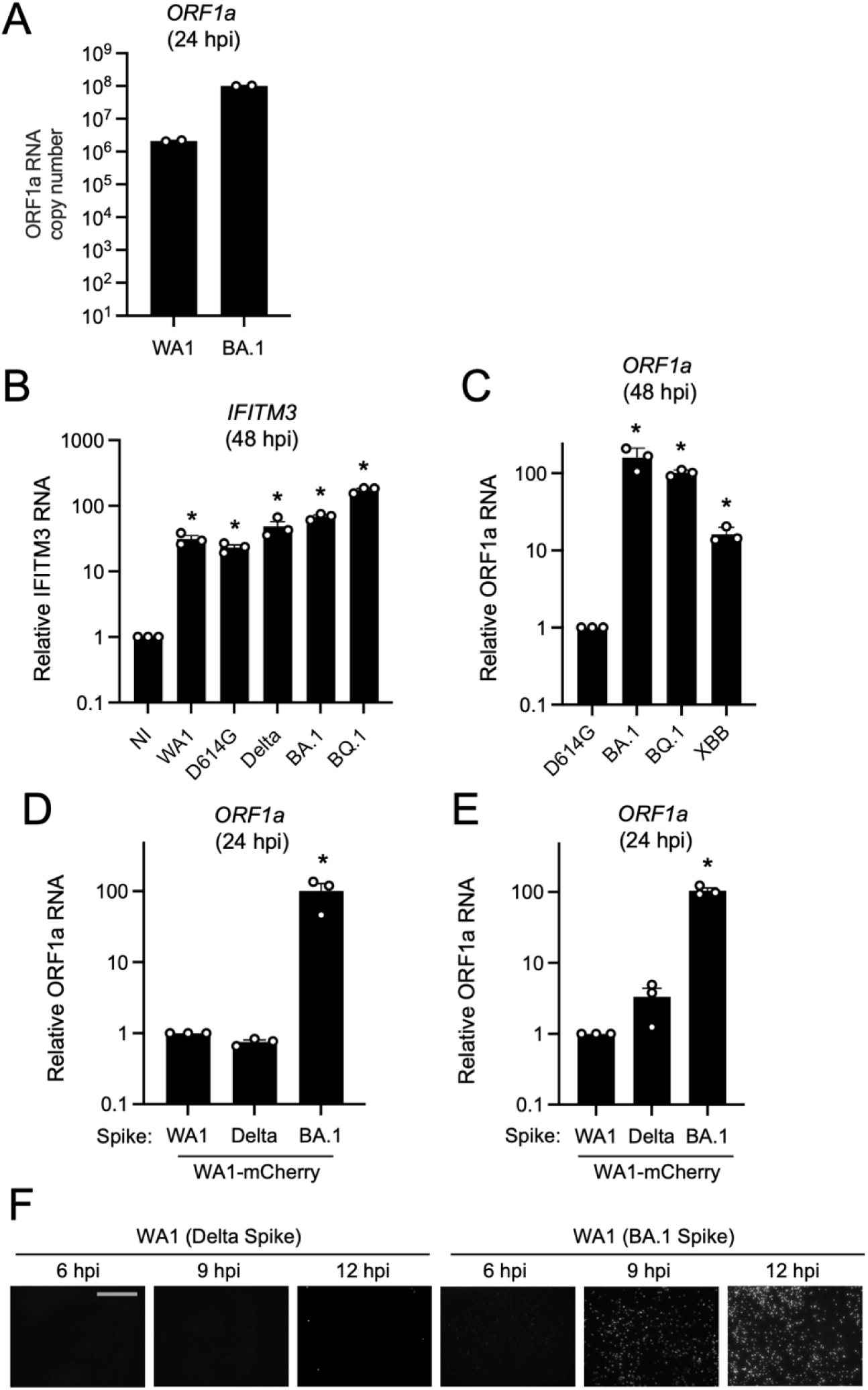
(A) Primary human nasal epithelial cells (pooled from 14 human donors) were cultured at the air-liquid interface and inoculated with 10000 plaque forming units of WA1 or BA.1. Total cellular RNA was extracted and viral ORF1a was quantified by RT-qPCR at 24 hours post-inoculation. Absolute ORF1a RNA copy numbers were calculated by comparison to an ORF1a standard curve. (B) Primary human nasal epithelial cells (pooled from 14 human donors) were cultured at the air-liquid interface and inoculated with 10000 plaque forming units of WA1, D614G, Delta, BA.1, or BQ.1. RT-qPCR of cellular *IFITM3* was performed at 48 hours post inoculation. Relative *IFITM3* transcript abundance was compared to actin using the 2^(-ΔΔCT) method. *IFITM3* abundance in non-inoculated (NI) cells was set to 1. Statistically significant differences (* *P* < 0.05) between the indicated condition and the NI condition were determined by one-way ANOVA. (C) Primary human nasal epithelial cells (pooled from 14 human donors) were cultured at the air-liquid interface and inoculated with 10000 plaque forming units of D614G, BA.1, BQ.1, or XBB. At 48 hours post-inoculation, total cellular RNA was extracted and viral ORF1a was quantified by RT-qPCR. Relative ORF1a abundance was determined by comparing to actin using the 2^(-ΔΔCT) method. Statistically significant differences (* *P* < 0.05) between the indicated condition and D614G were determined by one-way ANOVA. (D) 10000 plaque forming units of recombinant WA.1 encoding mCherry and Spike protein from WA1, Delta, or BA.1 (WA1-mCherry (WA1 Spike), WA1-mCherry (Delta Spike), and WA1-mCherry (BA.1 Spike)) were used to inoculate primary human nasal epithelial cells (pooled from 14 human donors) cultured at the air-liquid interface. At 48 hours post-inoculation, total cellular RNA was extracted and viral ORF1a was quantified by RT-qPCR. Relative ORF1a abundance was determined by comparing to actin using the 2^(-ΔΔCT) method. Statistically significant differences (* *P* < 0.05) between the indicated condition and WA1-mCherry (WA1 Spike) were determined by one-way ANOVA. (E) As in (D), except that 5×10^7 copies of absolute ORF1a RNA were used as input. At 48 hours post-inoculation, infection was measured by viral ORF1a RT-qPCR. Relative ORF1a abundance was determined by comparing to actin using the 2^(-ΔΔCT) method. Statistically significant differences (* *P* < 0.05) between the indicated condition WA.1 were determined by one-way ANOVA. All results are represented as means plus standard error from three independent infections (symbols represent biological replicates). (F) 10000 plaque forming units of recombinant WA.1 encoding mCherry and Spike protein from Delta or BA.1 (WA1-mCherry (Delta Spike), and WA1-mCherry (BA.1 Spike)) were used to inoculate primary human nasal epithelial cells (pooled from 14 human donors) cultured at the air-liquid interface. At 6, 9, and 12 hours post-inoculation, infection was measured by mCherry fluorescence. Scale bar = 300 microns.

**Supplemental Figure 3.**
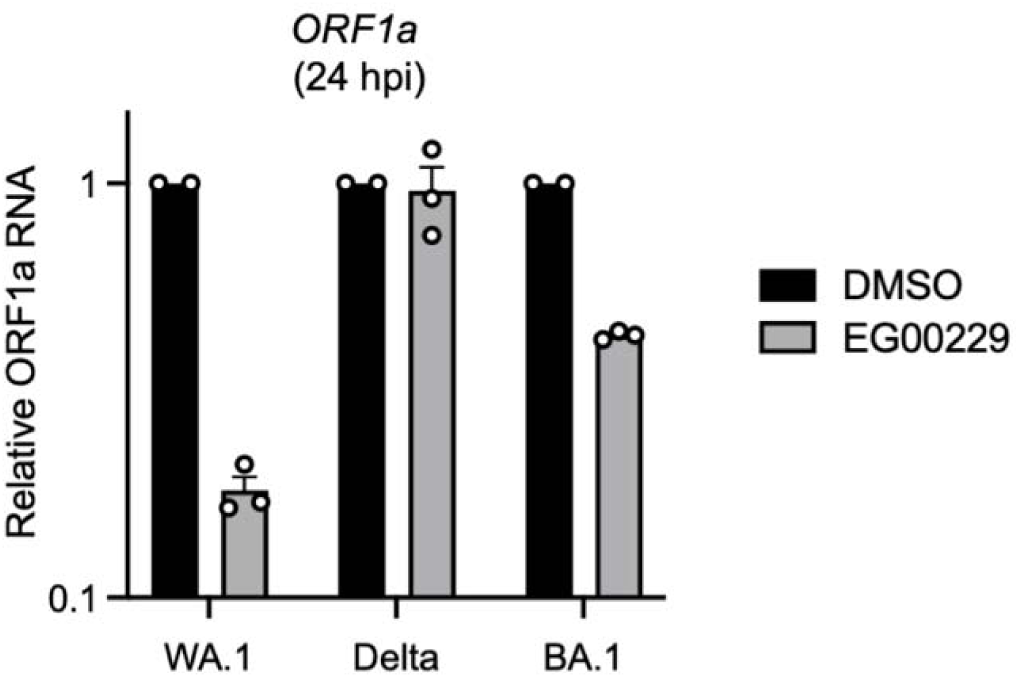
Primary human nasal epithelial cells (pooled from 14 human donors) were cultured at the air-liquid interface, pre-treated with 100 µM EG00229 or DMSO for two hours, and inoculated with 10000 plaque forming units of WA1, Delta, or BA.1. mCherry fluorescence was measured at 24 hours post-inoculation by high-content imaging. The fluorescence intensity of the DMSO-treated condition for each virus was set to 1. Scale bar = 300 µm. Results are represented as means plus standard error from one infection (symbols represent three RT-qPCR runs).

**Supplemental Figure 4.**
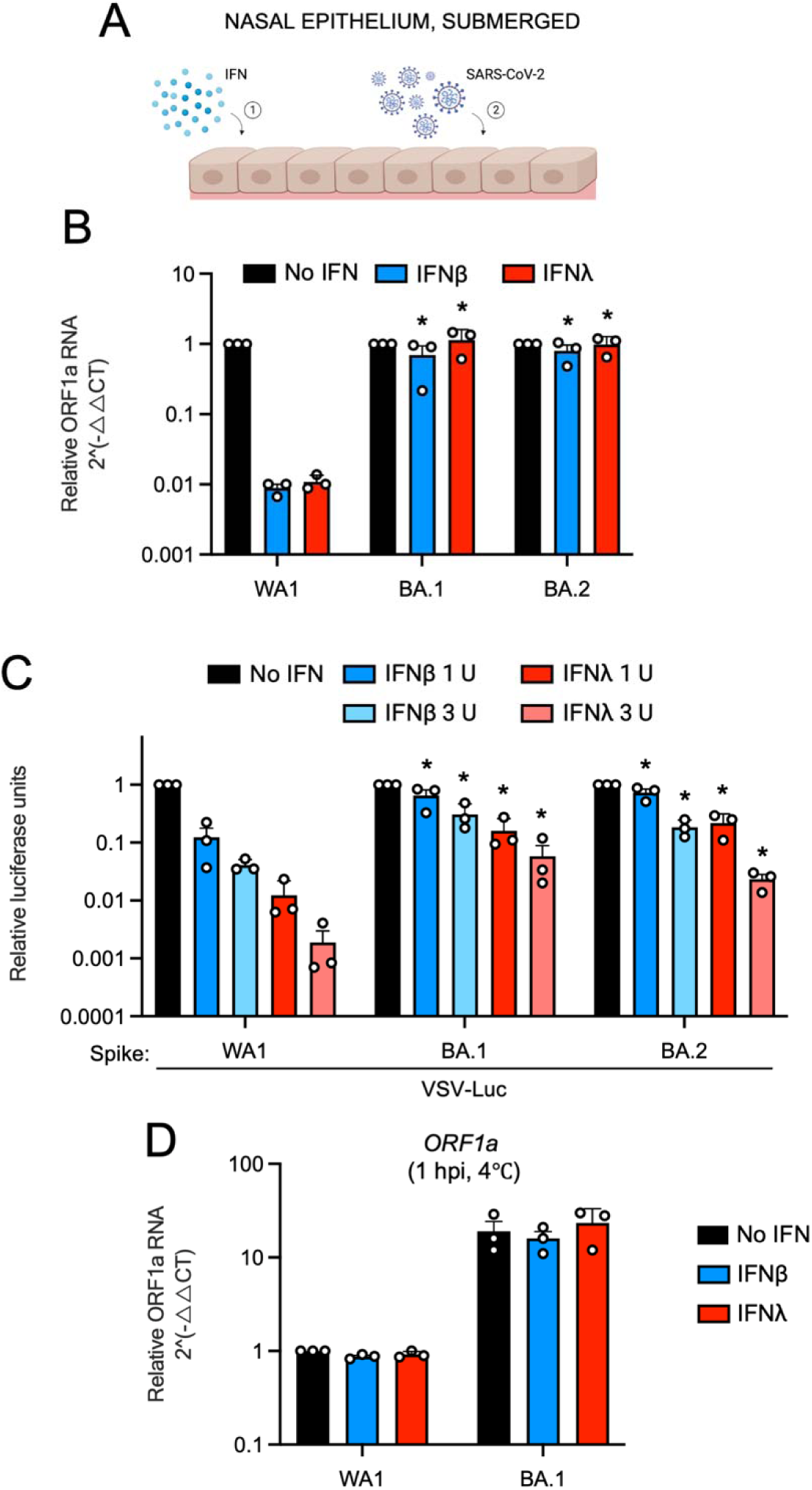
(A) Primary human nasal epithelial cells (pooled from 3 human donors) were cultured as undifferentiated, submerged monolayers, treated with IFN-beta or IFN-lambda for 18 hours, and challenged with SARS-CoV-2. Cartoon made with Biorender.com. (B) Cells were pre-treated with 2 units of IFN-beta or 2 units of FN-lambda for 18 hours, and inoculated with WA1, BA.1, or BA.2 at an MOI of 0.05. Total RNA was extracted from cells at 24 hours post inoculation, and ORF1a levels were measured by RT-qPCR. Relative ORF1a abundance was determined by comparing to actin using the 2^(-ΔΔCT) method. For each virus, ORF1a levels in the absence of IFN were set to 1. (C) Primary human nasal epithelial cells (pooled from 3 human donors) were pre-treated with the indicated amounts of IFN-beta or IFN-lambda for 18 hours and challenged with VSV-based pseudovirus decorated with Spike from WA1, BA.1, or BA.2. At 24 hours post inoculation, luciferase activity was measured from lysed cells. Luciferase activity of WA1, BA.1, and BA.2 pseudoviruses in the absence of IFN were set to 1. (D) Primary human nasal epithelial cells (pooled from 3 human donors) were pre-treated with 2 units of IFN-beta or 5 ng/mL IFN-lambda for 18 hours, and inoculated with WA1 or BA.1 at an MOI of 0.05 on ice. Total RNA was extracted from cells at 1 hour post inoculation, and ORF1a levels were measured by RT-qPCR. Relative ORF1a abundance was determined by comparing to actin using the 2^(-ΔΔCT) method. ORF1a levels of WA.1 in the absence of IFN were set to 1. All results are represented as means plus standard error from three independent infections (symbols represent biological replicates). Statistically significant differences (* *P* < 0.05) between the indicated condition and the corresponding No IFN condition were determined by one-way ANOVA.

**Supplemental Figure 5.**
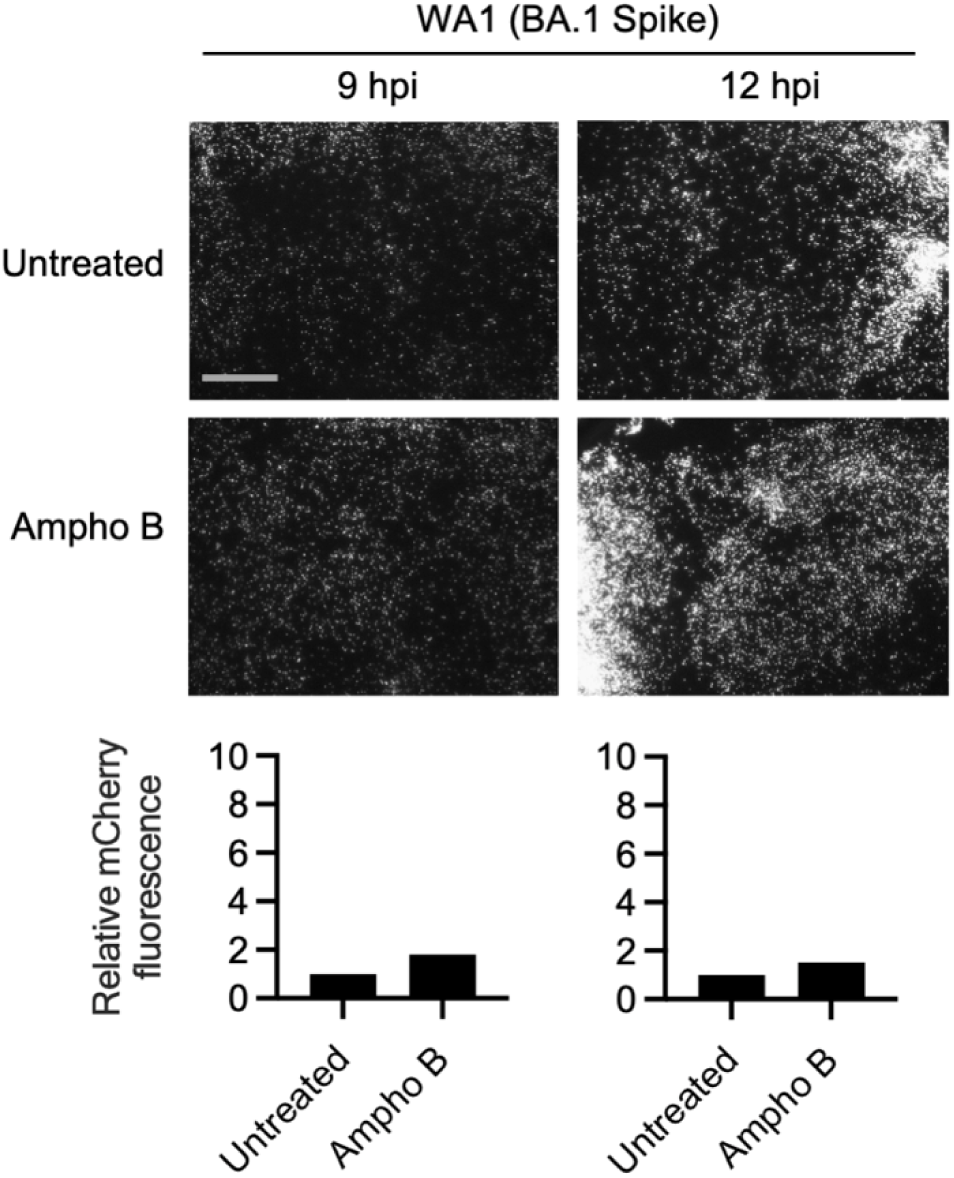
Primary human nasal epithelial cells (pooled from 14 human donors) were cultured at the air-liquid interface, pre-treated with 1 µM Amphotericin B for two hours or untreated, and inoculated with 10000 plaque forming units of WA1-mCherry (BA.1 Spike). mCherry fluorescence was measured at 9 and 12 hours post-inoculation by high-content imaging. The fluorescence intensity of the untreated condition was set to 1. Scale bar = 300 µm. Ampho B; amphotericin B.

## Materials and Methods

### Tissue culture

Frozen primary human nasal airway epithelial cells (hNAEC) were purchased from Epithelix and cultured as submerged monolayers with hAEC Culture Medium (Epithelix). hNAEC from three human donors were thawed and pooled, and cells were passed every 5 days. Fresh primary human nasal airway epithelial cells cultured at the liquid-air interface (nasal ALI) were obtained from Epithelix (MucilAir, Pool of Donors) and cultured with MucilAir culture medium (Epithelix) in trans-well plates provided by the company; culture medium was replaced every 2 days. Nasal ALI cultures were maintained for 2-5 weeks in our laboratories prior to inoculation with SARS-CoV-2. hNAEC and Nasal ALI were fixed with 4% paraformaldehyde, permeabilized with 0.1% Triton-X, and stained with anti-ACE2 (Abcam; ab15348), anti-Acetyl-a-Tubulin (Lys40) (D20G3) (Cell Signaling; 5335), anti-phalloidin (Cell Signaling; 8878), and DAPI to assess ACE2 levels, the presence of ciliated cells, and pseudostratification of nuclei, respectively. Images were obtained using the Zeiss LSM 880 workstation and AiryScan capability and were processed into 3D reconstructions by volume rendering using Imaris software. Primary human small airway (lung) epithelial cells (hSAEC) were purchased from ATCC (PCS-301-010) and were cultured with Airway Epithelial Cell Basal Medium (ATCC, PCS-300-030) and the Bronchial Epithelial Growth Kit (ATCC, PCS-300-400). hSAEC from three human donors were thawed and pooled, and cells were passed every 5 days. Vero E6 were purchased from ATCC (CRL-1586) and Vero E6-TMPRSS2 cells were generated from them by introducing the TMPRSS2 open reading frame using the Sleeping Beauty transposon system. Vero E6-ACE2-TMPRSS2 cells were obtained from BEI Resources (NR-54970). HEK293T-ACE2 cells were obtained from BEI Resources (NR-52511).

### Production of SARS-CoV-2 variants and infections

SARS-CoV-2 isolate USA/WA1/2020 (WA1; NR-52281), isolate hCoV-19/USA/HI-CDC-4359259-001/2021 (Lineage B.1.1.529; Omicron BA.1 variant; NR-56475), and isolate hCoV-19/USA/NY-MSHSPSP-PV56475/2022 (Lineage BA.2.12.1; Omicron BA.2 variant; NR-56782), isolate hCoV-19/USA/CA-Stanford-109_S21/2022 (Lineage XBB; Omicron XBB variant; NR-58927) and hCoV-19/USA/MDHP38960/2022 (Lineage BQ; Omicron BQ.1 variant) were obtained from BEI Resources. hCoV-19/New York-PV08410/2020 (Lineage B.1; D614G variant; NR-53514) was a gift from John Dye (USAMRIID). Isolate hCoV-19/USA/MD-HP05285/2021 VOC Delta G/478K.V1 (Lineage B.1.617.2 + AY.1 + AY.2; Delta variant) was obtained from Andrew Pekosz (Johns Hopkins University). Recombinant WA1-mCherry virus was rescued from a SARS-CoV-2 cDNA construct^75^, and mCherry was inserted at the N-terminus of the *N* gene. A P2A linker was placed between mCherry and N. To generate WA1-mCherry (BA.1 Spike) and WA1-mCherry (Delta Spike), the sequence encoding WA1 Spike was replaced with Spike from BA.1 or Delta, respectively. All viruses were grown in Vero E6-TMPRSS2 cells and harvested cell culture supernatants were titrated in Vero E6-TMPRSS2 cells to calculate infectious titers. Infectious titers were determined by focus-forming unit assay whereby foci of infection were measured using anti-nucleocapsid (N) antibody (Invitrogen; MA1-7403). Where indicated, ORF1a RNA titers were also calculated for some viruses to approximate virus particle quantity.

Infections of hNAEC or hSAEC monolayers were performed at a multiplicity of infection (MOI) of 0.05 as follows: cells were inoculated with virus suspension for 2 hours at 37°C; inoculum was removed; cells were washed with PBS and returned to 37°C for the time indicated. For detection of virus attachment, hNAEC were inoculated with virus suspension for 1 hour on ice; inoculum was removed; cells were washed three times with cold PBS.

For detection of infection or virus attachment by RT-qPCR, cells were lysed with Trizol (Sigma). Viral replication was measured using RT-qPCR amplification of viral ORF1a, as previously described^50^. Cells lysed with Trizol were mixed with chloroform (Sigma) at a 5:1 (Trizol:chloroform) ratio. Mixed samples were mixed thoroughly and incubated at room temperature for 10 minutes, followed by centrifugation at 12000 x G for 5 minutes to allow separation of the aqueous and organic phases. Equal volumes of 70% ethanol were added to the aqueous phases, mixed thoroughly, and incubated at room temperature for 5 minutes. RNA purification was performed using the PureLink RNA Mini Kit (Invitrogen) according to manufacturer’s instructions. Purified RNA product was immediately used with the One-step PrimeScript RT-PCR Kit (Takara). Primers and probes were obtained from IDT. The primers and probes used to amplify and quantify ORF1a are as follows (5’-3’): ORF1a-F AGAAGATTGGTTAGATGATGATAGT; ORF1a-R TTCCATCTCTAATTGAGGTTGAACC; ORF1a-P FAM/TCCTCACTGCCGTCTTGTTGACCA/BHQ13. The primers and probes used to amplify and quantify beta actin (ACTB) are as follows (5’-3’): ACTB-F ACAGAGCCTCGCCTTTG; ACTB-R CCTTGCACATGCCGGAG; ACTB-P 56-FAM/TCATCCATG/ZEN/GTGAGCTGGCGG/31ABkFQ. The primers and probes used to amplify and quantify IFITM3 are as follows (5’-3’): IFITM3-F ACCATGAATCACACTGTCCAAACCTT; IFITM3-R CCAGCACAGCCACCTCG; IFITM3-P FAM/ZEN-CTCTCCTGTCAACAGTGGCCAGCCCC-IBFQ. The primers and probes used to amplify and quantify IFNB are as follows (5’-3’): IFNB-F GAAACTGAAGATCTCCTAGCCT; IFNB-R GCCATCAGTCACTTAAACAGC; IFNB-P 56-FAM/TGAAGCAAT/ZEN/TGTCCAGTCCCAGAGG/3IABkFQ. Reaction mixtures of 20 µL (including 2.2 µL total RNA, 0.2 µM forward and reverse primers, and 0.1 µM probe) were subjected to reverse transcription (5 min at 45°C, followed by 10 s at 95°C) and 40 cycles of PCR (5 s at 95°C followed by 20 s at 60°C) in a CFX Opus 96 Real-Time PCR System (BioRad). Results were analyzed by the Comparative Ct Method (ΔΔCt Method) ^76^. RNA levels for viral ORF1a, IFNB, and IFITM3 were normalized to cellular ACTB.

For infections of primary nasal epithelial cells cultured at the air-liquid interface, the apical (air-exposed) surface was gently rinsed with 100 µL cell culture medium to partially remove mucous layers. Virus suspension in a volume of 50 µL was added to the apical surface and cells were incubated for 2 hours at 37°C. Inoculum was then removed and 100 µL PBS was used to gently wash the apical surface. Cells were then returned to 37°C. At 48 hours post inoculation, cells were lysed with Trizol and subjected to RNA extraction and RT-qPCR as indicated above. To quantify the infectious virus particles produced into the cell culture supernatant, medium was removed, 200 µL PBS was added to the apical surface, and cells were incubated at room temperature for 20 minutes before the PBS was recovered. A focus-forming units assay was performed by inoculating Vero E6-TMPRSS2 cells with the recovered volume. At 7 hours post inoculation, Vero E6-TMPRSS2 cells were fixed with 4% paraformaldehyde and stained with anti-N (Invitrogen; MA1-7403). The number of fluorescent foci was measured using a Cytation 5 Cell Imaging Multimode Reader (BioTeK).

For detection of recombinant WA-mCherry infection, the Cytation 5 Cell Imaging Multimode Reader (BioTeK) was used to measure mCherry fluorescence at the indicated hours post inoculation. Quantification of mCherry fluorescence was performed by measuring integrated signal intensities in Fiji.

### Production of VSV-Spike pseudoviruses and infections

Spike sequences from WA.1, BA.1, or BA.2 were codon-optimized for expression in human cells, synthesized with a 6xHis tag on the amino-terminus, and cloned into pcDNA3.1 (+) by GenScript. HEK293T cells were seeded in a 10 cm dish and transfected with 12 μg pcDNA3.1 Spike plasmid using Lipofectamine 2000 (Thermo Fisher). Forty-eight hours after transfection, culture medium was removed from cells, and 1 mL VSV-luc/GFP plus VSV-G (seed particles) was added. Twenty-four hours after infection, virus supernatants were collected, clarified by centrifugation at 500*g* for 5 minutes followed by filtration with a 45 μm filter, and stored at - 80°C. A total of 50 μL virus supernatants was added to submerged hNAEC from three pooled human donors, and 24 hours post inoculation, cells were lysed with Passive Lysis Buffer (Promega). Luciferase activity was measured on a Perkin Elmer MicroBeta 2450 microplate luminometer using the Luciferase Assay System (Promega). 50 μL volumes of VSV-WA1, VSV-BA.1, and VSV-BA.2 were found to infect Vero E6-TMPRSS2 cells to similar extents, suggestive of similar infectious titers.

### Interferons and Inhibitors

Recombinant human IFN-β (beta) (300-02BC) and human IFN-λ (lambda) (300-02L) were obtained from PeproTech and were used to test IFN sensitivity of full-length, authentic SARS-CoV-2 variants (IFN was added 18 hours prior to inoculation and removed prior to virus addition). Recombinant human IFN-β (beta) 1a (11415-1) and human IFN-λ (lambda) (11725-1) were obtained from PBL Assay Science and were used to test IFN sensitivity of VSV-based pseudoviruses (IFN was added 18 hours prior to inoculation and removed prior to virus addition). The following inhibitors were reconstituted in DMSO: E64d (Sigma; E8640), camostat mesylate (Sigma; SML0057), incyclinide (MedChemExpress; HY-13648), apratastat (MedChemExpress; HY-119307), batimastat (SelleckChem: S7155), EG00229 (SelleckChem; E1119). Bafilomycin A1 (Sigma; SML1661) was received as a ready-made solution in DMSO and amphotericin B (Sigma; A2942) was received as a ready-made solution in deionized water.

### Biosafety Approval

This study was conducted in compliance with all relevant local, state, and federal regulations. Approval for the generation and use of recombinant SARS-CoV-2 WA1-mCherry and its variants using Biosafety Level 3 practices was provided by the NIAID Institutional Biosafety Committee following evaluation by the Dual Use Research of Concern Institutional Review Entity (case number RD-22-X1-11; PI: Jonathan W Yewdell).

## Disclaimer

The content of this publication does not necessarily reflect the views or policies of the Department of Health and Human Services, nor does mention of trade names, commercial products, or organizations imply endorsement by the US Government.

## Acknowledgements

We would like to thank Brett Eaton, Elena Postnikova, Michael Murphy, Sushma Bhosle, Julie Tran, and Jillian Geiger (NIAID, Integrated Research Facility) for reagents and protocols to quantify viral ORF1a by RT-qPCR, Vincent Munster (NIAID) and Michael Letko (Washington State University) for reagents and protocols to produce VSV-Spike pseudoviruses, and Kim Peifley (NCI) for assistance in confocal immunofluorescence microscopy. Work in the laboratory of AAC was funded by the Intramural Research Program, NIH, NCI, Center for Cancer Research, and an Intramural Targeted Anti-COVID-19 award from NIAID. Work in the laboratory of JWY was funded by the Division of Intramural Research, NIH, NIAID.

## References

1 Letko, M., Marzi, A. & Munster, V. Functional assessment of cell entry and receptor usage for SARS-CoV-2 and other lineage B betacoronaviruses. Nature Microbiology 11, 1–17 (2020). 10.1038/s41564-020-0688-y

2 Hoffmann, M. et al. SARS-CoV-2 Cell Entry Depends on ACE2 and TMPRSS2 and Is Blocked by a Clinically Proven Protease Inhibitor. Cell, 1–19 (2020). 10.1016/j.cell.2020.02.052

3 Chu, H. et al. Host and viral determinants for efficient SARS-CoV-2 infection of the human lung. Nat Commun 12, 134 (2021). 10.1038/s41467-020-20457-w

4 Clausen, T. M. et al. SARS-CoV-2 Infection Depends on Cellular Heparan Sulfate and ACE2. Cell 183, 1043–1057 e1015 (2020). 10.1016/j.cell.2020.09.033

5 Carlos, A. J. et al. The chaperone GRP78 is a host auxiliary factor for SARS-CoV-2 and GRP78 depleting antibody blocks viral entry and infection. J Biol Chem 296, 100759 (2021). 10.1016/j.jbc.2021.100759

6 Cantuti-Castelvetri, L. et al. Neuropilin-1 facilitates SARS-CoV-2 cell entry and infectivity. Science 370, 856-+ (2020). 10.1126/science.abd2985

7 Daly, J. L. et al. Neuropilin-1 is a host factor for SARS-CoV-2 infection. Science 370, 861–865 (2020). 10.1126/science.abd3072

8 Wei, C. et al. HDL-scavenger receptor B type 1 facilitates SARS-CoV-2 entry. Nat Metab 2, 1391–1400 (2020). 10.1038/s42255-020-00324-0

9 Nguyen, L. et al. Sialic acid-containing glycolipids mediate binding and viral entry of SARS-CoV-2. Nat Chem Biol 18, 81–90 (2022). 10.1038/s41589-021-00924-1

10 Koch, J., Uckeley, Z. M., Doldan, P., Stanifer, M., Boulant, S. & Lozach, P. Y. TMPRSS2 expression dictates the entry route used by SARS-CoV-2 to infect host cells. EMBO J 40, e107821 (2021). 10.15252/embj.2021107821

11 Chan, J. F. et al. Altered host protease determinants for SARS-CoV-2 Omicron. Sci Adv 9, eadd3867 (2023). 10.1126/sciadv.add3867

12 Zang, R. et al. TMPRSS2 and TMPRSS4 promote SARS-CoV-2 infection of human small intestinal enterocytes. Sci Immunol 5 (2020). 10.1126/sciimmunol.abc3582

13 Laporte, M. et al. The SARS-CoV-2 and other human coronavirus spike proteins are fine-tuned towards temperature and proteases of the human airways. PLoS Pathog 17, e1009500 (2021). 10.1371/journal.ppat.1009500

14 Kishimoto, M. et al. TMPRSS11D and TMPRSS13 Activate the SARS-CoV-2 Spike Protein. Viruses 13 (2021). 10.3390/v13030384

15 V’Kovski, P., Kratzel, A., Steiner, S., Stalder, H. & Thiel, V. Coronavirus biology and replication: implications for SARS-CoV-2. Nat Rev Microbiol 19, 155–170 (2021). 10.1038/s41579-020-00468-6

16 Karim, S. S. A. & Karim, Q. A. Omicron SARS-CoV-2 variant: a new chapter in the COVID-19 pandemic. Lancet 398, 2126–2128 (2021). 10.1016/S0140-6736(21)02758-6

17 Pulliam, J. R. C. et al. Increased risk of SARS-CoV-2 reinfection associated with emergence of Omicron in South Africa. Science 376, eabn4947 (2022). 10.1126/science.abn4947

18 Dejnirattisai, W. et al. SARS-CoV-2 Omicron-B.1.1.529 leads to widespread escape from neutralizing antibody responses. Cell 185, 467-484 e415 (2022). 10.1016/j.cell.2021.12.046

19 Tuekprakhon, A. et al. Antibody escape of SARS-CoV-2 Omicron BA.4 and BA.5 from vaccine and BA.1 serum. Cell 185, 2422-2433 e2413 (2022). 10.1016/j.cell.2022.06.005

20 Nutalai, R. et al. Potent cross-reactive antibodies following Omicron breakthrough in vaccinees. Cell 185, 2116–2131 e2118 (2022). 10.1016/j.cell.2022.05.014

21 VanBlargan, L. A. et al. An infectious SARS-CoV-2 B.1.1.529 Omicron virus escapes neutralization by therapeutic monoclonal antibodies. Nat Med 28, 490-495 (2022). 10.1038/s41591-021-01678-y

22 Planas, D. et al. Considerable escape of SARS-CoV-2 Omicron to antibody neutralization. Nature 602, 671–675 (2022). 10.1038/s41586-021-04389-z

23 Wang, Q. et al. Antibody evasion by SARS-CoV-2 Omicron subvariants BA.2.12.1, BA.4 and BA.5. Nature 608, 603-608 (2022). 10.1038/s41586-022-05053-w

24 Lubin, J. H. et al. Structural models of SARS-CoV-2 Omicron variant in complex with ACE2 receptor or antibodies suggest altered binding interfaces. bioRxiv (2021). 10.1101/2021.12.12.472313

25 Huo, J. et al. A delicate balance between antibody evasion and ACE2 affinity for Omicron BA.2.75. Cell Rep 42, 111903 (2023). 10.1016/j.celrep.2022.111903

26 Jiang, X. L. et al. Omicron BQ.1 and BQ.1.1 escape neutralisation by omicron subvariant breakthrough infection. Lancet Infect Dis 23, 28-30 (2023). 10.1016/S1473-3099(22)00805-2

27 Cao, Y. et al. BA.2.12.1, BA.4 and BA.5 escape antibodies elicited by Omicron infection. Nature 608, 593-602 (2022). 10.1038/s41586-022-04980-y

28 Hong, Q. et al. Molecular basis of receptor binding and antibody neutralization of Omicron. Nature 604, 546–552 (2022). 10.1038/s41586-022-04581-9

29 Han, P. et al. Receptor binding and complex structures of human ACE2 to spike RBD from omicron and delta SARS-CoV-2. Cell 185, 630–640 e610 (2022). 10.1016/j.cell.2022.01.001

30 Peacock, T. P. et al. The altered entry pathway and antigenic distance of the SARS-CoV-2 Omicron variant map to separate domains of spike protein. bioRxiv, 2021.2012.2031.474653 (2022). 10.1101/2021.12.31.474653

31 Willett, B. J. et al. SARS-CoV-2 Omicron is an immune escape variant with an altered cell entry pathway. Nat Microbiol 7, 1161–1179 (2022). 10.1038/s41564-022-01143-7

32 Meng, B. et al. Altered TMPRSS2 usage by SARS-CoV-2 Omicron impacts infectivity and fusogenicity. Nature 603, 706–714 (2022). 10.1038/s41586-022-04474-x

33 Hui, K. P. Y. et al. SARS-CoV-2 Omicron variant replication in human bronchus and lung ex vivo. Nature 603, 715–720 (2022). 10.1038/s41586-022-04479-6

34 Qu, P. et al. Determinants and Mechanisms of the Low Fusogenicity and High Dependence on Endosomal Entry of Omicron Subvariants. mBio 14, e0317622 (2023). 10.1128/mbio.03176-22

35 Yuan, S. et al. Pathogenicity, transmissibility, and fitness of SARS-CoV-2 Omicron in Syrian hamsters. Science 377, 428–433 (2022). 10.1126/science.abn8939

36 Suzuki, R. et al. Attenuated fusogenicity and pathogenicity of SARS-CoV-2 Omicron variant. Nature 603, 700–705 (2022). 10.1038/s41586-022-04462-1

37 Zhao, H. et al. SARS-CoV-2 Omicron variant shows less efficient replication and fusion activity when compared with Delta variant in TMPRSS2-expressed cells. Emerg Microbes Infect 11, 277–283 (2022). 10.1080/22221751.2021.2023329

38 Halfmann, P. J. et al. SARS-CoV-2 Omicron virus causes attenuated disease in mice and hamsters. Nature 603, 687–692 (2022). 10.1038/s41586-022-04441-6

39 Kozlov, M. Omicron’s feeble attack on the lungs could make it less dangerous. Nature 601, 177 (2022). 10.1038/d41586-022-00007-8

40 Maslo, C., Friedland, R., Toubkin, M., Laubscher, A., Akaloo, T. & Kama, B. Characteristics and Outcomes of Hospitalized Patients in South Africa During the COVID-19 Omicron Wave Compared With Previous Waves. JAMA 327, 583–584 (2022). 10.1001/jama.2021.24868

41 Sposito, B. et al. The interferon landscape along the respiratory tract impacts the severity of COVID-19. Cell 184, 4953–4968 e4916 (2021). 10.1016/j.cell.2021.08.016

42 Reggio, A. et al. Role of FAM134 paralogues in endoplasmic reticulum remodeling, ER-phagy, and Collagen quality control. EMBO reports 22, e52289 (2021). 10.15252/embr.202052289

43 Wu, C. T. et al. SARS-CoV-2 replication in airway epithelia requires motile cilia and microvillar reprogramming. Cell 186, 112–130 e120 (2023). 10.1016/j.cell.2022.11.030

44 Bojkova, D., Widera, M., Ciesek, S., Wass, M. N., Michaelis, M. & Cinatl, J., Jr. Reduced interferon antagonism but similar drug sensitivity in Omicron variant compared to Delta variant of SARS-CoV-2 isolates. Cell Res 32, 319–321 (2022). 10.1038/s41422-022-00619-9

45 Jocher, G. et al. ADAM10 and ADAM17 promote SARS-CoV-2 cell entry and spike protein-mediated lung cell fusion. EMBO Rep 23, e54305 (2022). 10.15252/embr.202154305

46 Guo, K. et al. Interferon resistance of emerging SARS-CoV-2 variants. Proc Natl Acad Sci U S A 119, e2203760119 (2022). 10.1073/pnas.2203760119

47 Majdoul, S. & Compton, A. A. Lessons in self-defence: inhibition of virus entry by intrinsic immunity. Nature Reviews Immunology (2021). 10.1038/s41577-021-00626-8

48 Shi, G., Schwartz, O. & Compton, A. A. More than meets the I: the diverse antiviral and cellular functions of interferon-induced transmembrane proteins. Retrovirology 14, 1–11 (2017). 10.1186/s12977-017-0377-y

49 Kenney, A. D. et al. Interferon-induced transmembrane protein 3 (IFITM3) limits lethality of SARS-CoV-2 in mice. EMBO Rep 24, e56660 (2023). 10.15252/embr.202256660

50 Shi, G. et al. Rapalogs downmodulate intrinsic immunity and promote cell entry of SARS-CoV-2. J Clin Invest 132 (2022). 10.1172/JCI160766

51 Shi, G. et al. Opposing activities of IFITM proteins in SARSLCoVL2 infection. The EMBO journal 3, e201900542–201900512 (2020). 10.15252/embj.2020106501

52 Winstone, H. et al. The polybasic cleavage site in the SARS-CoV-2 spike modulates viral sensitivity to Type I interferon and IFITM2. Journal of Virology (2021). 10.1128/JVI.02422-20

53 Prelli Bozzo, C., et al. IFITM proteins promote SARS-CoV-2 infection and are targets for virus inhibition in vitro. Nat Commun 12, 4584 (2021). 10.1038/s41467-021-24817-y

54 Lin, T.-Y. et al. Amphotericin B Increases Influenza A Virus Infection by Preventing IFITM3-Mediated Restriction. Cell Reports 5, 895–908 (2013). 10.1016/j.celrep.2013.10.033

55 Wrensch, F. et al. Interferon-Induced Transmembrane Protein-Mediated Inhibition of Host Cell Entry of Ebolaviruses. The Journal of Infectious Diseases, jiv255 (2015). 10.1093/infdis/jiv255

56 Suddala, K. C. et al. Interferon-induced transmembrane protein 3 blocks fusion of sensitive but not resistant viruses by partitioning into virus-carrying endosomes. PLoS Pathogens 15, e1007532–1007535 (2019). 10.1371/journal.ppat.1007532

57 Rahman, K., Coomer, C. A., Majdoul, S., Ding, S. Y., Padilla-Parra, S. & Compton, A. A. Homology-guided identification of a conserved motif linking the antiviral functions of IFITM3 to its oligomeric state. eLife 9, 975–925 (2020). 10.7554/eLife.58537

58 Huotari, J. & Helenius, A. Endosome maturation. EMBO J 30, 3481–3500 (2011). 10.1038/emboj.2011.286

59 Nie, C., Sahoo, A. K., Netz, R. R., Herrmann, A., Ballauff, M. & Haag, R. Charge Matters: Mutations in Omicron Variant Favor Binding to Cells. Chembiochem 23, e202100681 (2022). 10.1002/cbic.202100681

60 Shi, D. et al. Structural Characteristics of Heparin Binding to SARS-CoV-2 Spike Protein RBD of Omicron Sub-Lineages BA.2.12.1, BA.4 and BA.5. Viruses 14 (2022). 10.3390/v14122696

61 Gelbach, A. L. et al. Interactions between heparin and SARS-CoV-2 spike glycoprotein RBD from omicron and other variants. Front Mol Biosci 9, 912887 (2022). 10.3389/fmolb.2022.912887

62 Baindara, P., Roy, D., Mandal, S. M. & Schrum, A. G. Conservation and Enhanced Binding of SARS-CoV-2 Omicron Spike Protein to Coreceptor Neuropilin-1 Predicted by Docking Analysis. Infect Dis Rep 14, 243–249 (2022). 10.3390/idr14020029

63 Beyer, D. K. & Forero, A. Mechanisms of Antiviral Immune Evasion of SARS-CoV-2. J Mol Biol 434, 167265 (2022). 10.1016/j.jmb.2021.167265

64 Lei, X. et al. Activation and evasion of type I interferon responses by SARS-CoV-2. Nat Commun 11, 3810 (2020). 10.1038/s41467-020-17665-9

65 Xia, H. et al. Evasion of Type I Interferon by SARS-CoV-2. Cell Rep 33, 108234 (2020). 10.1016/j.celrep.2020.108234

66 Shalamova, L. et al. Omicron variant of SARS-CoV-2 exhibits an increased resilience to the antiviral type I interferon response. PNAS Nexus 1, pgac067 (2022). 10.1093/pnasnexus/pgac067

67 Bojkova, D. et al. Omicron-induced interferon signalling prevents influenza A virus infection. bioRxiv, 2022.2009.2006.506799 (2022). 10.1101/2022.09.06.506799

68 Singh, J. et al. BA.1, BA.2 and BA.2.75 variants show comparable replication kinetics, reduced impact on epithelial barrier and elicit cross-neutralizing antibodies. PLoS Pathog 19, e1011196 (2023). 10.1371/journal.ppat.1011196

69 Alfi, O. et al. SARS-CoV-2 Omicron Induces Enhanced Mucosal Interferon Response Compared to other Variants of Concern, Associated with Restricted Replication in Human Lung Tissues. Viruses 14 (2022). 10.3390/v14071583

70 Prokunina-Olsson, L. et al. COVID-19 and emerging viral infections: The case for interferon lambda. J Exp Med 217 (2020). 10.1084/jem.20200653

71 Chong, Z. et al. Nasally delivered interferon-lambda protects mice against infection by SARS-CoV-2 variants including Omicron. Cell Rep 39, 110799 (2022). 10.1016/j.celrep.2022.110799

72 Jagannathan, P. et al. Peginterferon Lambda-1a for treatment of outpatients with uncomplicated COVID-19: a randomized placebo-controlled trial. Nat Commun 12, 1967 (2021). 10.1038/s41467-021-22177-1

73 Feld, J. J. et al. Peginterferon lambda for the treatment of outpatients with COVID-19: a phase 2, placebo-controlled randomised trial. Lancet Respir Med 9, 498–510 (2021). 10.1016/S2213-2600(20)30566-X

74 Reis, G. et al. Early Treatment with Pegylated Interferon Lambda for Covid-19. N Engl J Med 388, 518–528 (2023). 10.1056/NEJMoa2209760

75 Ye, C. et al. Rescue of SARS-CoV-2 from a Single Bacterial Artificial Chromosome. mBio 11 (2020). 10.1128/mBio.02168-20

76 Schmittgen, T. D. & Livak, K. J. Analyzing real-time PCR data by the comparative C(T) method. Nat Protoc 3, 1101–1108 (2008). 10.1038/nprot.2008.73

